# Cross-regulation between RpfG and HtsHs to regulate antibiotic biosynthesis in *Lysobacter*

**DOI:** 10.1101/2020.07.13.201541

**Authors:** Kaihuai Li, Gaoge Xu, Bo Wang, Guichun Wu, Fengquan Liu

## Abstract

Bacterial two-component systems (TCSs) sense and respond to environmental changes and modulate downstream gene expression. However, the mechanism of cross-talk between multiple TCSs is unclear. In this study, we report a previously uncharacterized mechanism by which the TCS protein RpfG interacts with hybrid two-component system (HyTCS) proteins HtsH1, HtsH2 and HtsH3 to regulate antibiotic biosynthesis in *Lysobacter*. RpfG, a phosphodiesterase (PDE), can degrade c-di-GMP to 5’-pGpG and can regulate antibiotic heat-stable antifungal factor (HSAF) biosynthesis in a PDE- independent manner. Thus, we wondered whether RpfG regulate HSAF biosynthesis through interactions with other factors. Subsequently, we demonstrated that RpfG interacts with three HyTCS proteins (HtsH1, HtsH2 and HtsH3), that can inhibit the PDE enzymatic activity of RpfG. Importantly, deletion of *htsH1*, *htsH2* and *htsH3* resulted in significantly decreased HSAF production, and we showed that HtsH1, HtsH2 and HtsH3 depend on their phosphorylation activity to directly regulate HSAF biosynthesis gene expression. Our results reveal that RpfG does not depend on PDE activity to regulate HSAF biosynthesis, rather it interacts with HtsH1, HtsH2 and HtsH3 to do so, a regulatory mechanism that may be a conserved paradigm in *Lysobacter* and *Xanthomonas*.

## Introduction

Signal transduction mediated by two-component systems (TCSs), including hybrid two-component systems (HyTCSs), is widely employed by bacteria to sense and respond to environmental changes. A typical two-component regulatory system is a signal transduction pathway consisting of a histidine protein kinase (HK) receptor and a response regulator (RR) (Azcarate-Peril *et al.*, 2005). Usually, as each particular TCS is specialized to respond to a specific environmental signal, cross-regulation between multiple TCSs may be present in a single bacterial cell (Goodman *et al.*, 2009). However, how cross-talk between multiple TCSs remains incompletely studied.

The diffusible signal factor (DSF) represents a new class of widely conserved quorum sensing (QS) signals with a cis-2-unsaturated fatty acid moiety that regulate various biological functions in pathogenic and beneficial environmental bacteria (Deng *et al.*, 2011; Qian *et al.*, 2013). The Rpf gene cluster is important for the DSF signalling network in bacteria, and the role of RpfF and RpfC/RpfG TCS in this gene cluster in DSF production and signal transduction has been well documented (Barber *et al.*, 1997; Chatterjee & Sonti, 2002; He *et al.*, 2006; Ryan *et al.*, 2010; Deng *et al.*, 2011). A previous study revealed that RpfG contains both an N-terminal response regulator domain and a C-terminal HD-GYP domain (Andrade *et al.*, 2006). HD-GYP domain proteins are cyclic dimeric GMP (c-di-GMP) phosphodiesterases (PDEs) that can degrade c-di-GMP (Bellini *et al.*, 2014). The HD-GYP of RpfG domain interacts with the hybrid histidine kinase RpfC to link QS and c-di-GMP signalling pathways in bacteria (Andrade *et al.*, 2006). However, whether RpfG can also interact with other TCSs to regulate different pathways remains unknown.

*L*. *enzymogenes* is a nonpathogenic strain that was used to control crop fungal diseases known for synthesis of the heat-stable antifungal factor (HSAF), which exhibits inhibitory activity against a wide range of fungal species (Zhao *et al.*, 2019; Zhao *et al.*, 2017; Qian *et al.*, 2009; Odhiambo *et al.*, 2017; Lou *et al.*, 2011; Yu *et al.*, 2007; Li *et al.*, 2006; Li *et al.*, 2018). Our earlier work revealed that RpfG affects HSAF production in *L. enzymogenes* (Xu *et al.*, 2018). However, the molecular mechanism by which RpfG regulates the biosynthesis of HSAF remains unknown. We discovered that RpfG interacts with three HyTCS proteins through bioinformatics predictions (Figure 3—figure supplement 1). We designated the HyTCS protein HtsH (<u>H</u>ybrid two-component <u>s</u>ignalling system regulating <u>H</u>SAF production) based on the findings of this study.

In the present study, we demonstrated that RpfG interacts with HtsHs to regulate the biosynthesis of the antifungal antibiotic HSAF. We describe for the first time the regulatory functions of HtsH1, HtsH2 and HtsH3 in bacteria. These functions likely represent a common mechanism that helps establish signalling specificity in bacteria possessing cross-talk among different TCSs.

## Results

### The HD-GYP domain of RpfG has PDE activity and can degrade c-di-GMP

Sequence analysis revealed that the HD-GYP domain contains all residues essential for PDE activities, thus suggesting that RpfG may be a PDE enzyme. To obtain direct evidence for the biochemical function of RpfG, recombinant N-terminal maltose binding protein (MBP) RpfG (designated RpfG-MBP) was produced. The proteins had a monomeric molecular weight of 71 kDa and were purified by dextrin sepharose high performance to obtain preparations that exhibited single bands by SDS-gel electrophoresis (Figure 1A). This RpfG-MBP protein was able to degrade the model substrate c-di-GMP to 5’-pGpG, consistent with its PDE activity (Figure 1B). Quantitative analysis revealed that RpfG-MBP exhibited a high level of activity for the degradation of c-di-GMP with 100% degraded at 5 min after initiation of the reaction in comparison with the MBP enzyme as a control (Figure 1C). To better understand the roles of the HD-GYP domain in RpfG function, we substituted the RpfG residues His-190, Asp-191, Gly-253, Tyr-254 and Pro-255 of the HD-GYP signature motif with alanine (Ala) by site-directed mutagenesis into constructs expressing the RpfG-H190A-MBP, RpfG-D191A-MBP, RpfG-G253A-MBP and RpfG-P255A-MBP proteins. We tested the c-di-GMP PDE activity of these mutant proteins. Our results showed that point mutations of the His-190, Asp-191, Gly-253 and Pro-255 residues in RpfG almost abolished PDE activity (Figure 1B, C). These data suggest that RpfG has PDE activity and that the HD-GYP domain is required for full PDE activity of RpfG in vivo.

**Figure 1.**
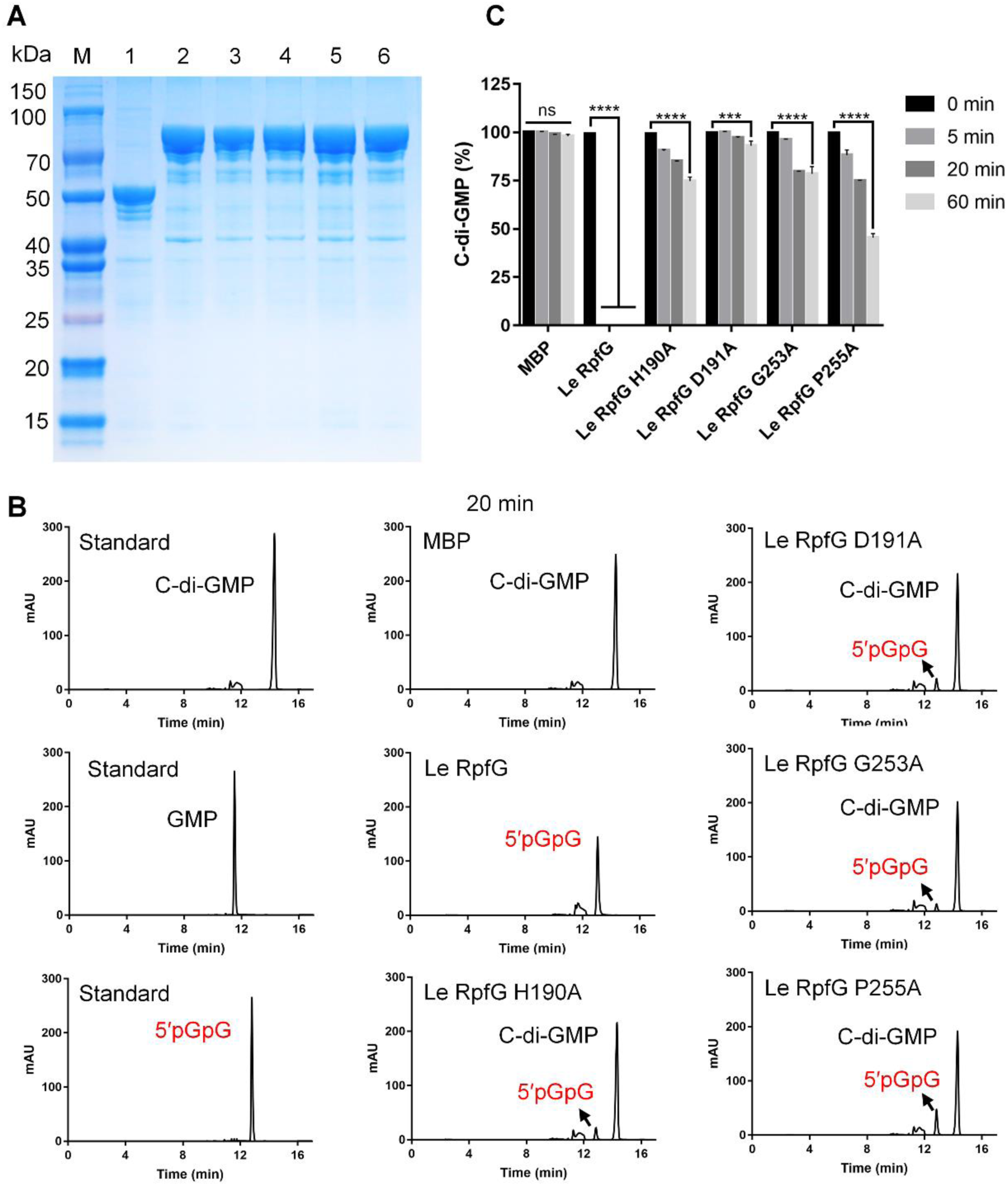
RpfG is a PDE enzyme. **(A)** The purified protein was analysed by 12% SDS-PAGE. Lane M, molecular mass markers; lane 1, MBP protein; lane 2, RpfG-MBP protein; lane 3, RpfG-MBP H190A protein; lane 4, RpfG-MBP D191A protein; lane 5, RpfG-MBP G253A protein; lane 6, RpfG-MBP P255A protein. **(B)** The purified protein has enzymatic activity against c-di-GMP. HPLC analysis of aliquots of reaction mixtures boiled immediately after addition of the enzyme and after 20 min of incubation shows the degradation of c-di-GMP to a compound with the same mobility as the 5′pGpG standard. **(C)** The purified RpfG protein has PDE enzymatic activity. For convenience of comparison, the peak of c-di-GMP in the RpfG solution at 0 min was defined as 100% and used to normalize the c-di-GMP level ratios at different time points. Error bars, means ± standard deviations (n = 3). ∗∗∗ P < 0.001, ∗∗∗∗ P < 0.0001, assessed by one-way ANOVA. All experiments were repeated three times with similar results.

### RpfG regulatory activity does not depend on its PDE enzymatic activity

Our earlier work revealed that RpfG affects HSAF production in *L. enzymogenes* (Xu *et al.*, 2018). However, the molecular mechanism by which RpfG regulates HSAF synthesis remains unknown. Since RpfG, as a PDE enzyme, was able to degrade the substrate c-di-GMP to 5′-pGpG, we asked whether RpfG could function to regulate HSAF biosynthesis depending on the PDE enzymatic activity in *L. enzymogenes*. We quantified HSAF production in the Δ*rpfG* mutant and complementary strain (Δ*rpfG*/*rpfG*) by HPLC. We found that HSAF production in the Δ*rpfG* mutant strain was completely suppressed (Figure 2A). The complementary strain Δ*rpfG*/*rpfG* yielded HSAF at the level of the wild-type strain (Figure 2A). These results were similar to those of the above research (Xu *et al.*, 2018). To examine the relationship between the regulatory and enzymatic activities of RpfG, mutations at the conserved His-190, Asp-191, Gly-253, Tyr-254 and Pro-255 of the HD-GYP signature motif with alanine (Ala) were examined by site-directed mutagenesis. We tested HSAF production in the Δ*rpfG* mutant strain carrying plasmids encoding these mutant proteins. The strains expressing the RpfG H190A, D191A, G253A, Y254A and P255A mutant proteins resulted in increased HSAF production compared with that of the Δ*rpfG* mutant strain. These results were similar to those of the complementary strain Δ*rpfG*/*rpfG*. Importantly, the Δ*rpfG* mutant strain and complemented strains (Δ*rpfG*/*rpfG*, Δ*rpfG*/*rpfG* H190A, Δ*rpfG*/*rpfG* D191A, Δ*rpfG*/*rpfG* G253A, Δ*rpfG*/*rpfG* Y254A and Δ*rpfG*/*rpfG* P255A) did not impair bacterial growth (Figure 2B, C), implying that Le RpfG plays a specific role in regulating HSAF production. In previous work, His-190, Asp-191, Gly-253, Tyr-254 and Pro-255 were found to be critical for the PDE activity of RpfG (Figure 1); therefore, we expected that RpfG regulated HSAF synthesis independent of PDE enzymatic activity. To test this prediction, we compared intracellular c-di-GMP concentrations in the Δ*rpfG* mutant and the wild type and in the HSAF-production medium (1/10 TSB). We found that the concentration in the Δ*rpfG* mutant did not significantly change c-di-GMP production compared to that in the wild-type strain (Figure 2D). These findings indicated that the regulatory activity of RpfG does not depend on its PDE enzymatic activity against c-di-GMP.s

**Figure 2.**
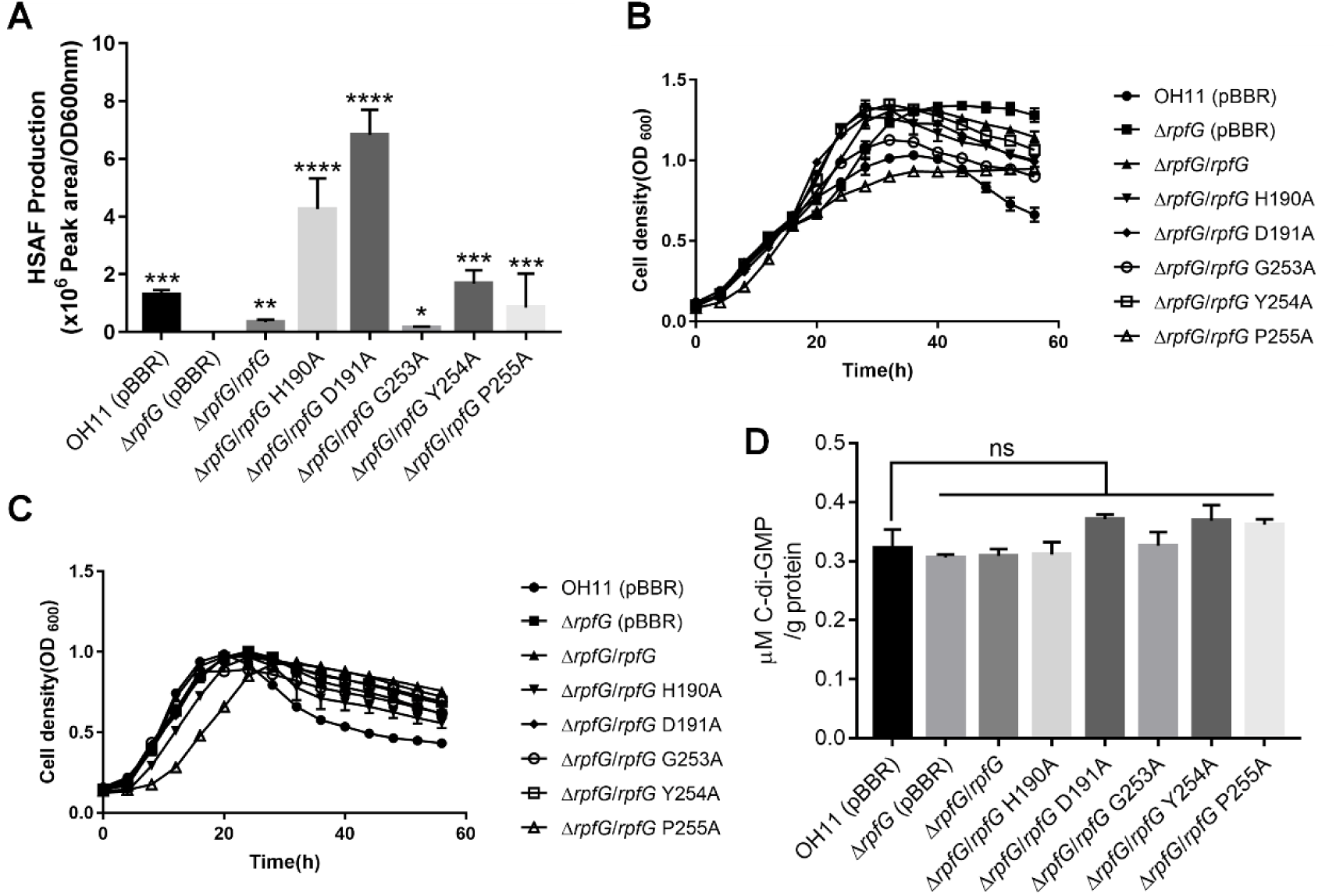
RpfG-mediated regulation of HSAF production does not depend on its PDE activity. **(A)** Quantification of HSAF produced by the *rpfG* mutant strain and strains complemented with *rpfG* or the *rpfG* site-directed mutant genes grown in 10% TSB medium. **(B)** Growth curves of the *rpfG* mutant strain and strains complemented with *rpfG* or the *rpfG* site-directed mutant genes in rich LB medium. **(C)** Growth curves of the *rpfG* mutant strain and strains complemented with the *rpfG* or *rpfG* site-directed mutant genes in 10% TSB medium. **(D)** Intracellular c-di-GMP concentrations in the wild type, *rpfG* mutant and strains complemented with *rpfG* or the *rpfG* site-directed mutant genes. Error bars, means ± standard deviations (n = 3). ∗ P < 0.05, ∗∗ P < 0.01, ∗∗∗ P < 0.001, ∗ ∗∗∗ P < 0.0001, assessed by one-way ANOVA. All experiments were repeated three times with similar results.

### RpfG binds directly to HyTCS protein HtsHs

Previous studies confirmed that RpfG does not regulate HSAF biosynthesis through the c-di-GMP signalling pathway, indicating that RpfG may regulate HSAF biosynthesis through interactions with other proteins in *L. enzymogenes*. To further explore the mechanisms underlying the contribution of RpfG to HSAF production, we performed bioinformatics predictions. We found that RpfG may interact with three HyTCS proteins (HtsH1, HtsH2 and HtsH3) (Figure 3—figure supplement 1A). To verify the operon structure of *htsHs* for in-depth genetic analyses, a series of RT-PCR primers (Supplementary Table S2 and Figure 3—figure supplement 1B) were designed to determine whether there are intergenic transcripts crossing the adjacent genes. As shown in Figure 3—figure supplement 1C, Le3071 (*htsH1*), Le3072 (*htsH2*) and Le3073 (*htsH3*) likely constitute a single transcription unit because the corresponding intergenic transcripts were successfully amplified. Le3071 (*htsH1*), Le3072 (*htsH2*) and Le3073 (*htsH3*) encode a group of typical HyTCS proteins with pfam Reg_prop, pfam Y-Y-Y, HisKA, HATPase_c, and Response_reg domains. All three HyTCS proteins contain one predicted transmembrane region (Figure 3—figure supplement 1D). We examined the alignments of three HyTCS proteins (HtsH1, HtsH2 and HtsH3). The results showed that the HtsH1 protein shares 50% and 53% identity with HtsH2 and HtsH3, respectively. We also aligned HtsH2 with HtsH3, and the identity values were 50% (Figure 3—figure supplement 2).

To examine the possibility that RpfG binds directly to the HyTCS proteins (HtsH1, HtsH2 and HtsH3), we used a pull-down assay using *E. coli*-expressed proteins *in vitro*. We purified recombinant RpfG-MBP, and the cytoplasmic fragments of HtsH1, HtsH2 and HtsH3 (HtsH1C-Flag-His, HtsH2C-HA-His and HtsH3C-Myc-His, respectively) from *E. coli* (Figure 1A and Figure 3—figure supplement 3). First, we tested the ability of RpfG-MBP to pull down HtsH1C-Flag-His, HtsH2C-HA-His and HtsH3C-Myc-His. Finally, we confirmed the interaction between RpfG and HtsH1, HtsH2 or HtsH3 using pull down assays (Figure 3A). On the other hand, we examined the ability of HtsH1C-Flag-His, HtsH2C-HA-His and HtsH3C-Myc-His to pull down RpfG-MBP and observed a positive signals (Figure 3B-D). Second, we used surface plasmon resonance (SPR) to measure the possible binding events between the RpfG and HtsH1, HtsH2 or HtsH3 proteins. The RpfG-MBP sensor physically bound HtsH1C-Flag-His with a binding constant (K_D_) of 0.06675 nM (Figure 3E), suggesting an intermediate level of protein-protein interaction. We also confirmed direct binding between the RpfG-MBP and HtsH2C-HA-His or HtsH3C-Myc-His proteins by SPR (K_D_ = 0.2998 nM or K_D_ = 0.1678 nM, respectively) (Figure 3F-G). Additionally, the SPR assay revealed that HtsH1C-Flag-His bound to HtsH2C-HA-His or HtsH3C-Myc-His with reasonably high affinity (K_D_ = 0.09619 nM or K_D_ = 0.1597 nM, respectively) and revealed that HtsH2C-HA-His bound HtsH3C-Myc-His with a K_D_ value of 0.1782 nM (Figure 3—figure supplement 4). Taken together, these experiments demonstrate that RpfG directly interacts with HtsH1, HtsH2 or HtsH3 proteins *in vitro*.

**Figure 3.**
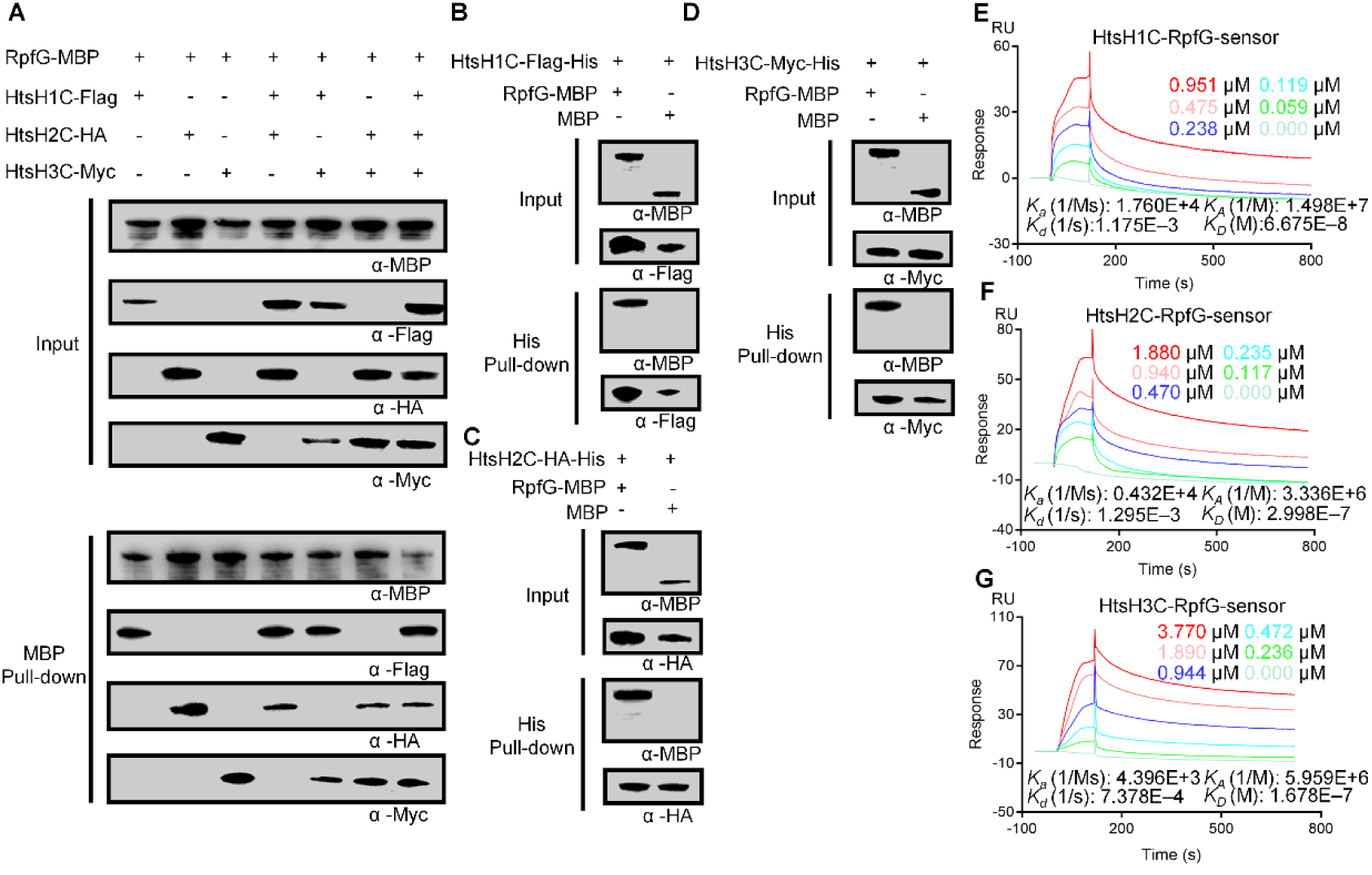
RpfG interaction with HtsH1, HtsH2 and HtsH3. **(A)** An MBP pull-down assay confirming interactions between RpfG-MBP and the cytoplasmic fragment of HtsH1, HtsH2 and HtsH3 (HtsH1C-Flag-His, HtsH2C-HA-His and HtsH3C-Myc-His, respectively). The pull-down assay was carried out using anti-MBP antibody. Western blotting was performed using anti-MBP, anti-Flag, anti-HA and anti-Myc antibodies. **(B)** A His pull-down assay confirming interactions between the cytoplasmic fragment of HtsH1 (HtsH1C-Flag-His) and RpfG-MBP. The pull-down assay was carried out using Ni-nitrilotriacetic acid (NTA) agarose. Western blotting was performed using anti-Flag and anti-MBP antibodies. **(C)** A His pull-down assay confirming interactions between the cytoplasmic fragment of HtsH2 (HtsH2C-HA-His) and RpfG-MBP. The pull-down assay was carried out using Ni-NTA agarose. Western blotting was performed using anti-HA and anti-MBP antibodies. **(D)** A His pull-down assay confirming interactions between the cytoplasmic fragment of HtsH3 (HtsH3C-Myc-His) and RpfG-MBP. The pull-down assay was carried out using Ni-NTA agarose. Western blotting was performed using anti-Myc and anti-MBP antibodies. **(E)** SPR showing that HtsH1C-Flag-His forms a complex with RpfG-MBP with K_D_ = 0.06675 nM. **(F)** SPR showing that HtsH2C-HA-His forms a complex with RpfG-MBP with K_D_ = 0.2998 nM. **(G)** SPR showing that HtsH3C-Myc-His forms a complex with RpfG-MBP with K_D_ = 0.1678 nM.

### HtsHs inhibit the PDE enzymatic activity of RpfG

We wondered whether RpfG and HtsH1, HtsH2 or HtsH3 interactions affect the PDE activity of RpfG. To test this possibility, we used a biochemical assay in which c-di-GMP hydrolysis by RpfG-MBP was assayed in the presence or absence of HtsH1, HtsH2 or HtsH3. The results of the assay showed that the PDE activity of RpfG-MBP was lower in the presence than in the absence of HtsH1C-Flag-His, HtsH2C-HA-His or HtsH3C-Myc-His (Figure 4). Therefore, the results of the assays suggested that HtsH1, HtsH2 and HtsH3 inhibit the PDE enzymatic activity of RpfG. This result further confirms that the ability of RpfG to regulate HSAF production does not depend on its PDE enzymatic activity against c-di-GMP in *L. enzymogenes*.

**Figure 4.**
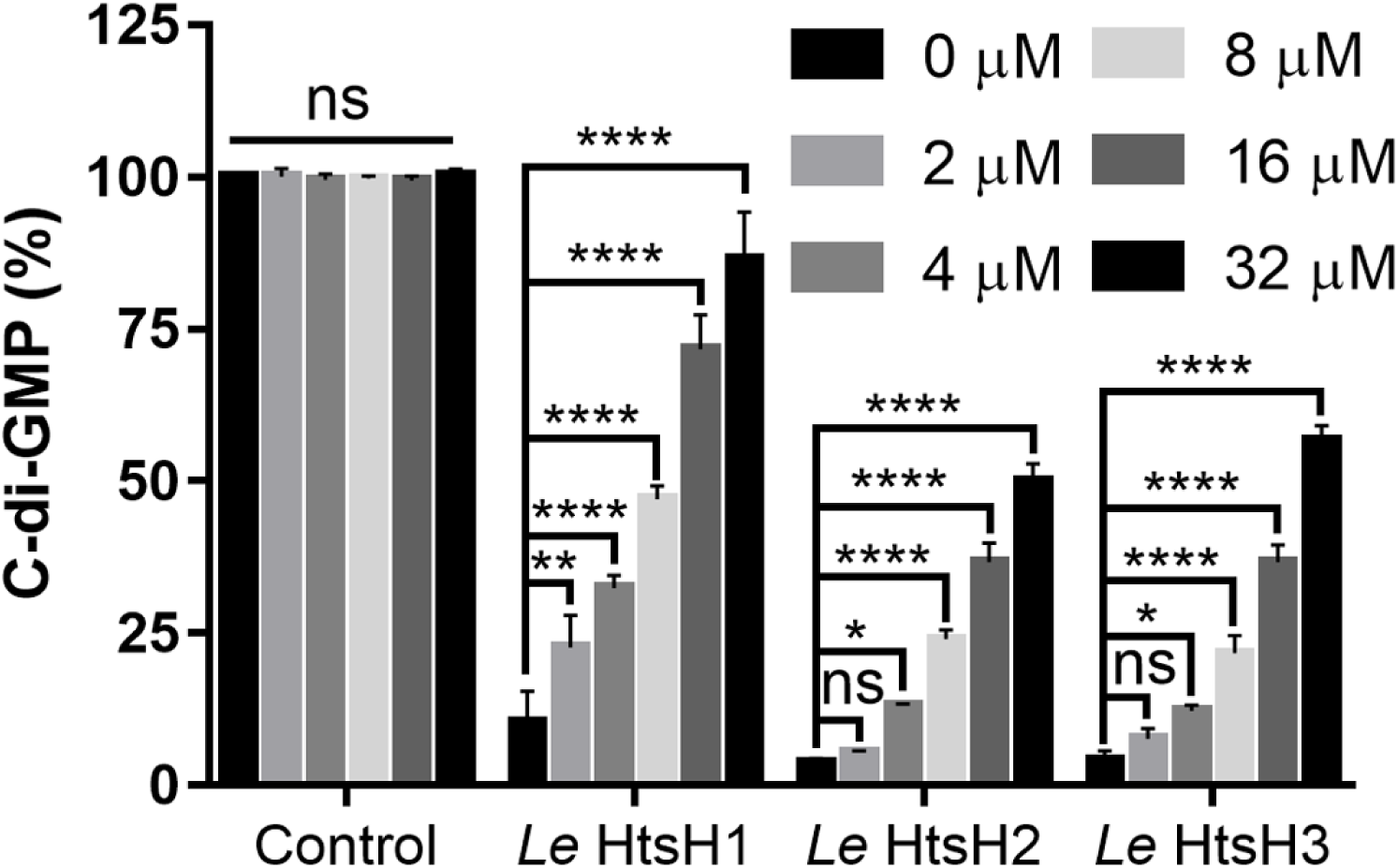
Impact of HtsH1, HtsH2 and HtsH3 on RpfG enzyme activity. The addition of HtsH1C-Flag-His, HtsH2C-HA-His and HtsH3C-Myc-His proteins to an RpfG-MBP protein solution decreased its c-di-GMP phosphodiesterase activity. For convenience of comparison, the peak of c-di-GMP in the MBP solution was defined as 100% and used to normalize the c-di-GMP level ratios at 20 min.

### Deletion of *htsH1*, *htsH2* and *htsH3* resulted in decreased HSAF production

The above studies found that RpfG may regulate HSAF biosynthesis through interaction with HtsHs (HtsH1, HtsH2 and HtsH3) in *L. enzymogenes*. To identify the physiological functions of HtsH1, HtsH2 and HtsH3 in HSAF production, the genes *htsH1* (Le3071), *htsH2* (Le3072) and *htsH3* (Le3073) were deleted using a two-step homologous recombination approach to construct the single knockout strains Δ*htsH1*, Δ*htsH2*, and Δ*htsH3*; the double mutant knockout strains Δ*htsH12*, Δ*htsH13*, and Δ*htsH23*; and triple knockout strain Δ*htsH123*. Subsequently, we quantified HSAF production in all of the mutant strains described above by HPLC. We found that Δ*htsH1*, Δ*htsH2* and Δ*htsH3* exhibited weakly decreased HSAF levels. However, the double-mutant strains Δ*htsH12*, Δ*htsH13*, Δ*htsH23* and triple-mutant strain Δ*htsH123* exhibited a significant decrease in HSAF levels compared with the wild-type levels (Figure 5A). To determine the role of HtsH1, HtsH2 and HtsH3 in the regulation of HSAF biosynthesis, we complemented Δ*htsH1*, Δ*htsH2*, Δ*htsH3* and the double-mutant strains Δ*htsH12*, Δ*htsH13*, Δ*htsH23* and triple-mutant strain Δ*htsH123* with plasmid-borne *htsH1*, *htsH2*, *htsH3*, *htsH12*, *htsH13*, *htsH23* and *htsH123*. HSAF production in the complemented strains (Δ*htsH1/htsH1*, Δ*htsH2*/*htsH2*, Δ*htsH3/htsH3*, Δ*htsH12/htsH12*, Δ*htsH13/htsH13*, Δ*htsH23/htsH23* and Δ*htsH123/htsH123*) restored HSAF biosynthesis compared with the wild-type levels (Figure 5A). Importantly, Δ*htsH1*, Δ*htsH2*, Δ*htsH3*, the double-mutant strains Δ*htsH12*, Δ*htsH13*, Δ*htsH23* and the triple-mutant strain Δ*htsH123* did not impair bacterial growth (Figure 5B, C), implying that HtsH1, HtsH2 and HtsH3 play a specific role in regulating HSAF production.

**Figure 5.**
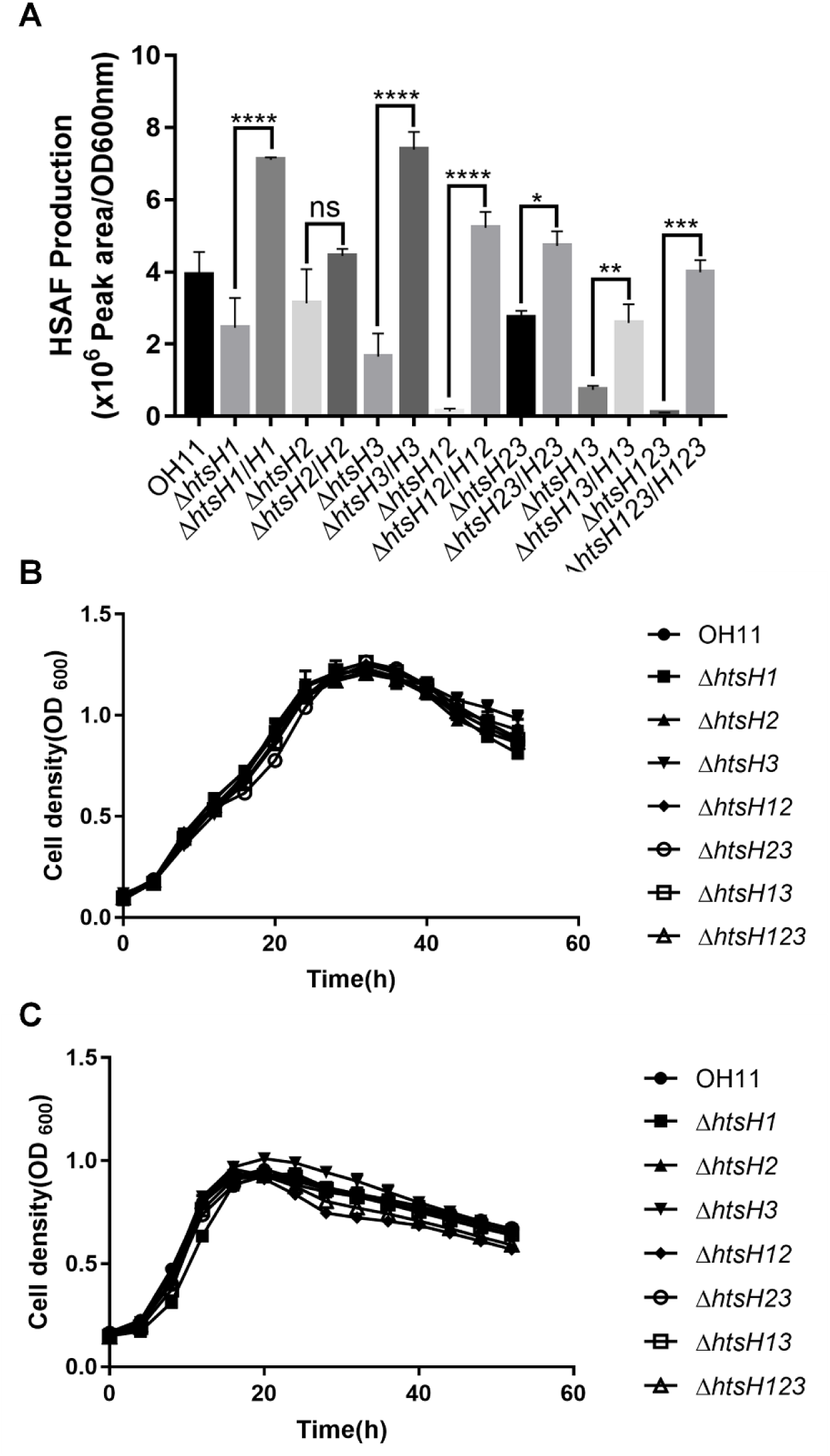
Deletion of *htsH1*, *htsH2* and *htsH3* resulted in decreased HSAF production. **(A)** Quantification of the HSAF produced by the *htsH1*, *htsH2* and *htsH3* mutant strains and strains complemented with the *htsH1*, *htsH2* and *htsH3* genes grown in 10% TSB medium. **(B)** Growth curves of the *htsH1*, *htsH2* and *htsH3* mutant strains and strains complemented with the *htsH1*, *htsH2* and *htsH3* genes in LB-rich medium. **(C)** Growth curves of the *htsH1*, *htsH2* and *htsH3* mutant strains and strains complemented with the *htsH1*, *htsH2* and *htsH3* genes in 10% TSB medium. Error bars, means ± standard deviations (n = 3). ∗ P < 0.05, ∗∗ P < 0.01, ∗∗∗ P < 0.001, ∗∗∗∗ P < 0.0001, assessed by one-way ANOVA. All experiments were repeated three times with similar results.

### HtsH1, HtsH2 and HtsH3 directly positively regulated HSAF biosynthesis gene expression

Earlier, we found that deletion of *htsH1*, *htsH2* and *htsH3* resulted in decreased HSAF production. We speculated that HtsH1, HtsH2 and HtsH3 might directly target HSAF biosynthesis gene promoters. To test this hypothesis, we performed an *E. coli*-based one-hybrid assay. As shown in Figure 6A, HtsH1, HtsH2 and HtsH3 could directly bind to the promoter of the HSAF biosynthesis gene (p*lafB*).

**Figure 6.**
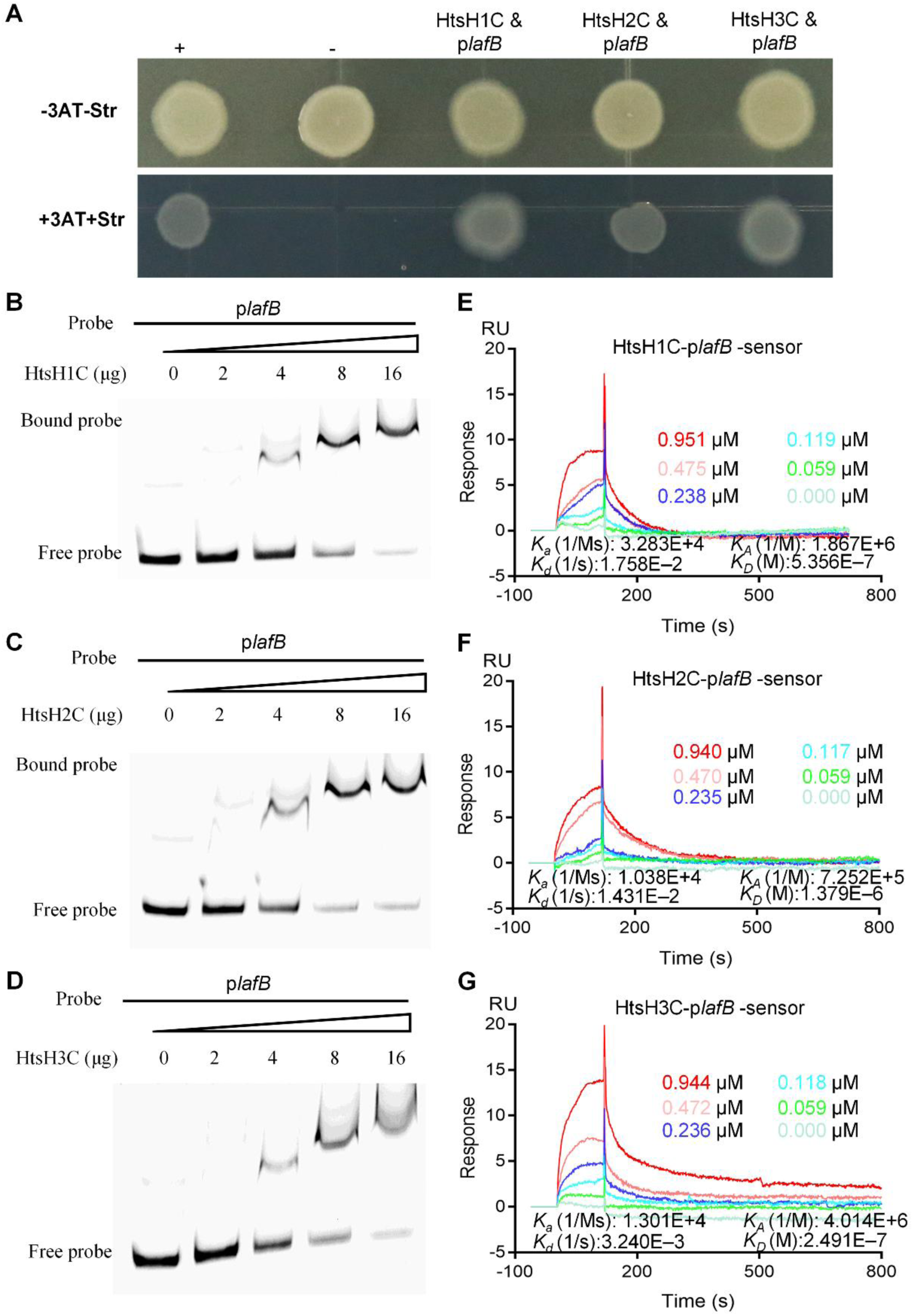
HtsH1, HtsH2 and HtsH3 directly bound the promoter of the HSAF biosynthesis gene (p*lafB*). **(A)** Direct physical interaction between HtsH1, HtsH2 and HtsH3 and the *lafB* promoter region was detected in *E. coli*. Experiments were performed according to the procedures described in the Materials and Methods section. +, cotransformant containing pBX-R2031 and pTRG-R3133, used as a positive control; -, cotransformant containing pBXcmT and the empty pTRG, used as a negative control; HtsH1C & p*lafB*, cotransformant harbouring pTRG-HtsH1C and pBXcmT-p*lafB*; HtsH2C & p*lafB*, cotransformant harbouring pTRG-HtsH2C and pBXcmT-p*lafB*; HtsH3C & p*lafB*, cotransformant harbouring pTRG-HtsH3C and pBXcmT-p*lafB*. -3AT-Str, plate with no selective medium (3AT, 3-amino-1,2,4-triazole and Str, streptomycin) and +3AT+Str, plate with M9-based selective medium **(B-D)** Gel shift assay showing that HtsH1, HtsH2 and HtsH3 directly regulate an HSAF biosynthesis gene. HtsH1C-Flag-His, HtsH2C-HA-His or HtsH3C-Myc-His protein (0, 1, 2, 4 or 8 μM) was added to reaction mixtures containing 50 ng of probe DNA, and the reaction mixtures were separated on polyacrylamide gels. **(E)** SPR showing that HtsH1C-Flag-His forms a complex with p*lafB* with K_D_ = 0.5356 nM. **(F)** SPR showing that HtsH2C-HA-His forms a complex with p*lafB* with K_D_ = 1.379 nM. **(G)** SPR showing that HtsH3C-Myc-His forms a complex with p*lafB* with K_D_ = 0.2491 nM.

Next, we tested the ability of HtsH1, HtsH2 and HtsH3 to bind to the *lafB* promoter, using an electrophoretic mobility shift assay (EMSA). A PCR-amplified 590 bp DNA fragment from the *lafB* promoter, namely, p*lafB*, was used as a probe. The addition of purified HtsH1C-Flag-His, HtsH2C-HA-His and HtsH3C-Myc-His protein, ranging from 0 µM to 10 µM, to the reaction mixtures (20 μL and at 28°C, 25 min) caused a shift in the mobility of the p*lafB* DNA fragment, and the EMSA revealed strong HtsH1, HtsH2 and HtsH3 binding with the p*lafB* probe in a dose-dependent manner (Figure 6B-D). We quantified the binding affinity of HtsH1, HtsH2 and HtsH3 to the HSAF operon promoter. In an SPR analysis, HtsH1C-Flag-His directly bound to the promoter of the HSAF biosynthesis gene (p*lafB*) with high affinity (K_D_ = 0.5356 nM) (Figure 6E). In addition, HtsH2C-HA-His and HtsH3C-Myc-His bound to p*lafB* with K_D_ values of 1.379 nM and 0.2491 nM, respectively (Figure 6F-G). The results demonstrated that HtsH1, HtsH2 and HtsH3 could directly target the promoters of the HSAF biosynthesis gene.

Based on the above results, we compared the transcriptome profiles of the wild-type strain and the *htsHs* mutants (Δ*htsH1*, Δ*htsH2*, Δ*htsH3*, Δ*htsH12*, Δ*htsH13*, Δ*htsH23* and Δ*htsH123*) by RNA-Seq and observed changes in the expression levels of several hundred genes (Supplementary Table S3). We then performed trend analysis of differential gene expression and found that the amounts of HSAF biosynthesis gene cluster mRNA were constitutively decreased in the *htsHs* mutants (Δ*htsH1*, Δ*htsH2*, Δ*htsH3*, Δ*htsH12*, Δ*htsH13*, Δ*htsH23* and Δ*htsH123*) (Figure 6—figure supplement 1).

Taken together, these results suggested that HtsH1, HtsH2 and HtsH3 can directly target the promoters of the HSAF biosynthesis genes to increase their expression and HSAF production by *L. enzymogenes*.

### HtsH1, HtsH2 and HtsH3 directly target HSAF biosynthesis gene promoters depending on their phosphorylation activity

Since HtsH1, HtsH2 and HtsH3 function as a group of HyTCS proteins, we explored whether HtsH1, HtsH2 and HtsH3 could directly target p*lafB* depending on its phosphorylation activity in *L. enzymogenes*. To this end, we assessed the phosphorylation levels of these proteins with calf intestine alkaline phosphatase (CIAP) in the reaction mixtures (20 μL, 28°C, 60 min). Using Mn^2+^-Phos-tag SDS-PAGE, we showed that the phosphorylation levels of HtsH1C-Flag-His, HtsH2C-HA-His and HtsH3C-Myc-His were decreased upon CIAP treatment (Figure 7A-C). To test the effect of HtsH1-, HtsH2- or HtsH3-mediated phosphorylation on the function of the target p*lafB*, EMSAs were performed with the same experimental conditions. The EMSA revealed that the amount of probes bound to HtsH1C-Flag-His, HtsH2C-HA-His and HtsH3C-Myc-His decreased with increasing amounts of CIAP (Figure 7D-F). The above findings indicated that HtsH1, HtsH2 and HtsH3 could directly regulate HSAF biosynthesis gene expression depending on their phosphorylation activity in *L. enzymogenes*.

**Figure 7.**
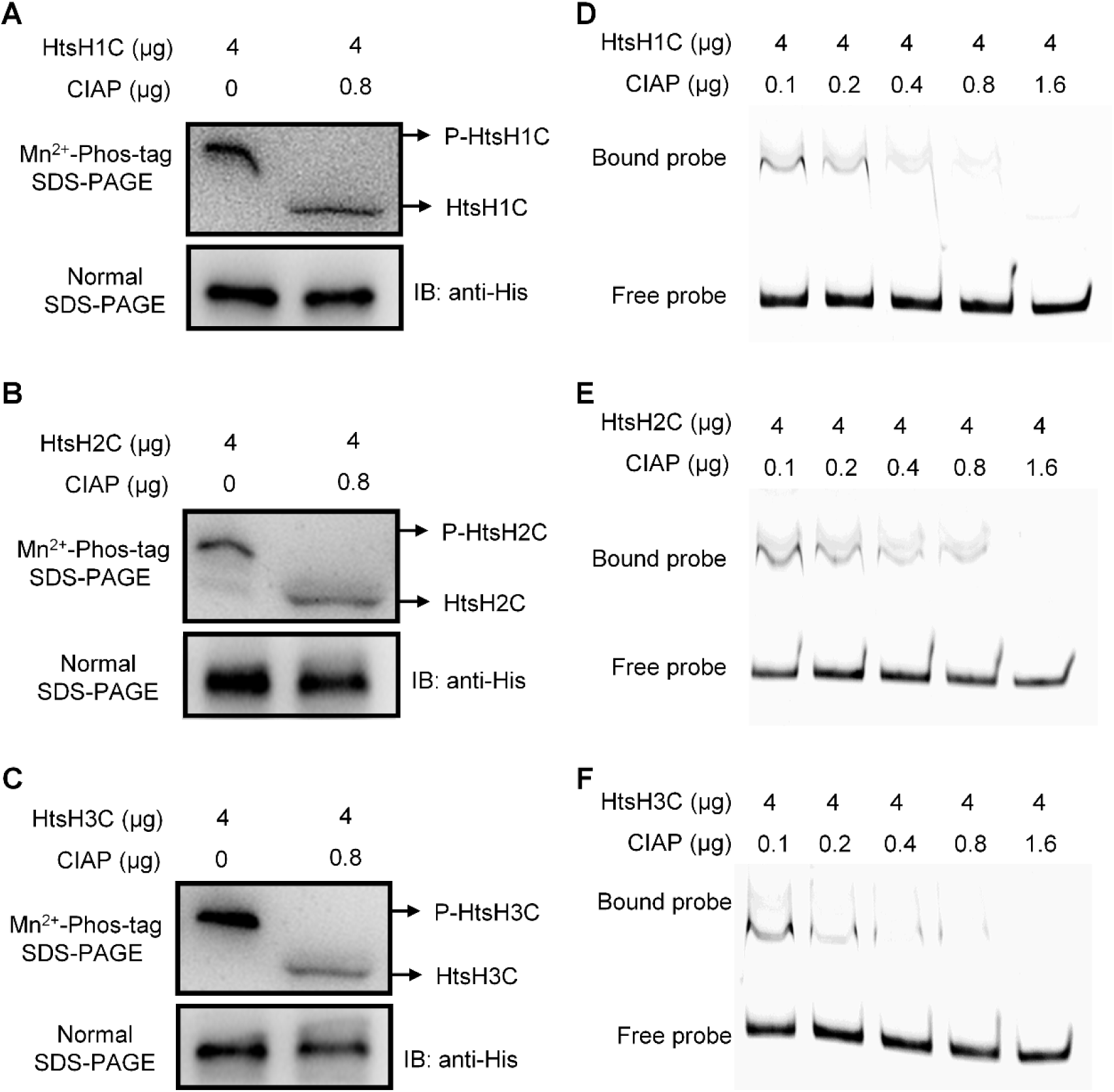
HtsH1, HtsH2 and HtsH3 directly target p*lafB* depending on their PDE activity. **(A-C)** Phosphorylation analysis of HtsH1C-Flag-His, HtsH2C-HA-His or HtsH3C-Myc-His with calf intestine alkaline phosphatase (CIAP) at concentrations ranging from 0.1 µg to 1.6 µg in the reaction mixtures (20 μL, 28°C, 60 min) using Mn^2+^- Phos-tag SDS-PAGE. **(D-F)** Gel shift assay showing that HtsH1C-Flag-His, HtsH2C-HA-His or HtsH3C-Myc-His does not directly target p*lafB* with calf intestine alkaline phosphatase (CIAP) at concentrations ranging from 0.1 µg to 1.6 µg in the reaction mixtures (20 μL, 28°C, 60 min). HtsHs (0, 1, 2, 4 or 8 μM) were added to reaction mixtures containing 50 ng of probe DNA, and the reaction mixtures were separated on polyacrylamide gels.

### RpfG- and HtsH1-, HtsH2- and HtsH3-dependent regulatory patterns are present in a wide range of bacterial species

In the present study, by using BLAST of the Nonredundant Protein Sequences (Nr) database of the National Center for Biotechnology Information (NCBI), we found that RpfG, the *rpf* cluster, and HtsH1, HtsH2 and HtsH3 were present not only in the genomes of *Lysobacter* species but also in *Xanthomonas* species (Figure 8—figure supplement 1). These findings suggest that RpfG- and HtsH1-, HtsH2- and HtsH3-dependent regulatory patterns are conserved mechanisms in *Lysobacter* and *Xanthomonas*.

## Discussion

In previous studies, we and our collaborators have shown that two-component systems (TCS) were employed by *L. enzymogenes* to affect HSAF production (Chen *et al.*, 2017; Su *et al.*, 2018; Zhou *et al.*, 2015). However, the mechanism through which TCS coordinates the synthesis of HSAF remains unknown. RpfG, as a member of the TCS family, contains a C-terminal HD-GYP domain that can affect HSAF production in *L. enzymogenes* (Andrade *et al.*, 2006; Han *et al.*, 2015). However, how RpfG regulates the synthesis of HSAF remains unknown. The results of the present study provides biochemical, genetic, and biophysical evidence to demonstrate for the first time that RpfG interacts with HtsHs to regulate the biosynthesis of the antifungal antibiotic HSAF. HtsH1, HtsH2 and HtsH3 regulate the expression of HSAF synthesis genes depending on their phosphorylation (Figure 8).

**Figure 8.**
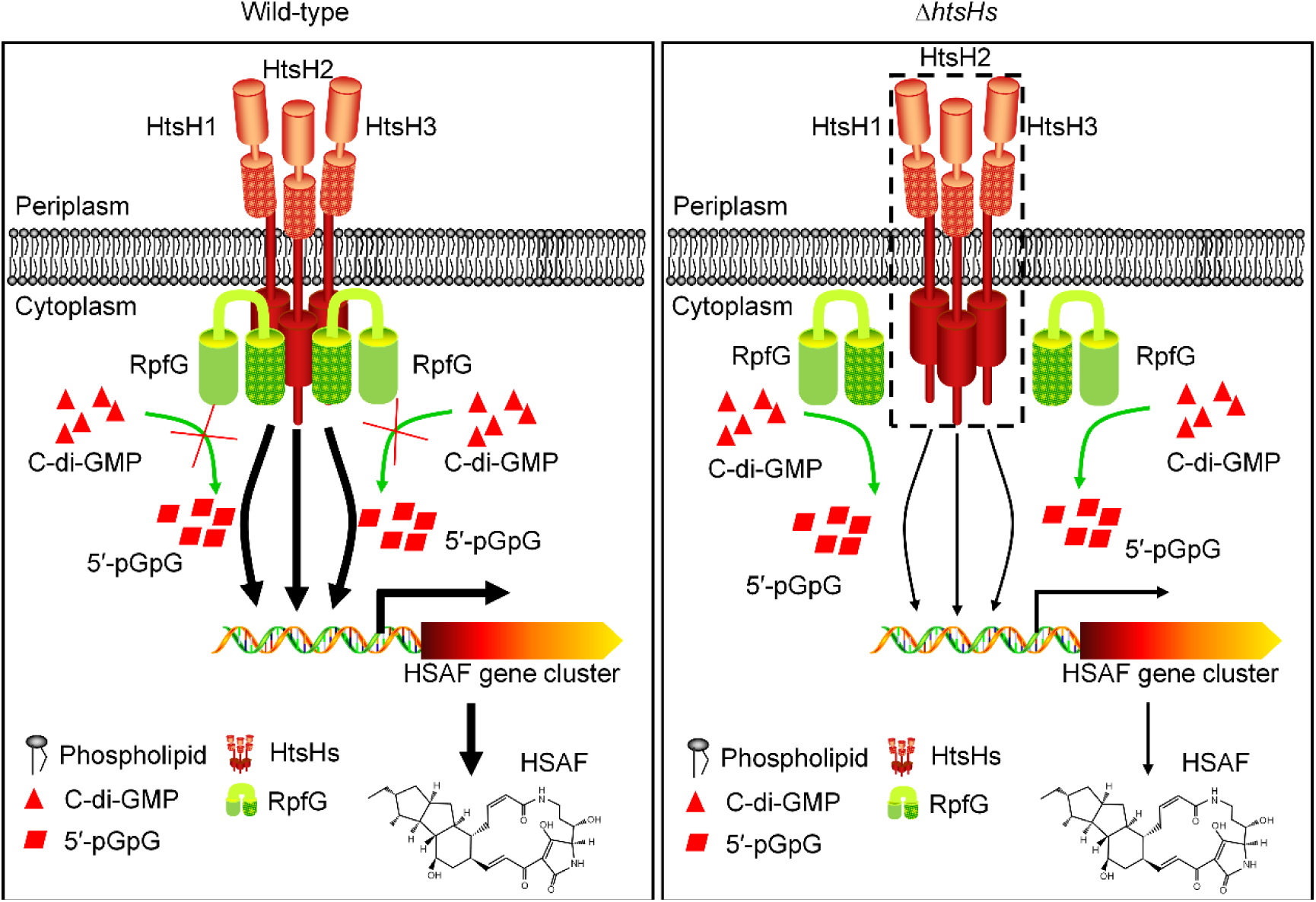
Schematic of the proposed RpfG directly interacting with three HyTCS proteins (HtsH1, HtsH2 and HtsH3) to regulate HSAF biosynthesis. The potential regulatory pathways and interactions of RpfG with HtsH1, HtsH2 and HtsH3 are proposed according to our observations and previous studies. RpfG and HtsH1, HtsH2 or HtsH3 interactions affect the PDE activity of RpfG. Phosphorylated HtsH1, HtsH2 and HtsH3 can directly target the promoter of HSAF biosynthesis genes to regulate HSAF production in *L. enzymogenes*.

HD-GYP domain proteins are c-di-GMP PDEs that can degrade c-di-GMP (Bellini *et al.*, 2014). However, the role of the HD-GYP domain of RpfG in the degradation of c-di-GMP in *L. enzymogenes* remained unelucidated. Therefore, we tested the PDE activity of RpfG and successfully showed that it was able to degrade the model substrate c-di-GMP to 5′-pGpG. To test whether the HD-GYP motif was important for catalytic activity in RpfG, we constructed RpfG mutant proteins (RpfG-H190A-MBP, RpfG-D191A-MBP, RpfG-G253A-MBP and RpfG-P255A-MBP). We tested the c-di-GMP PDE activity of these mutant proteins and suggested that the HD-GYP domain was required for full PDE activity of RpfG in vivo. It is generally speculated that the PDE activity of HD-GYP domain proteins is to degrade c-di-GMP to GMP (Galperin *et al.*, 1999; Galperin *et al.*, 2001; Andrade *et al.*, 2006). However, we showed for the first time that the activity of the HD-GYP domain of RpfG is involved in the degradation of c-di-GMP to 5′-pGpG.

In previous work, His-190, Asp-191, Gly-253, Tyr-254 and Pro-255 were found to be critical for the PDE activity of RpfG. Intriguingly, we found that the strains expressing the RpfG H190A, D191A, G253A, Y254A and P255A mutant proteins resulted in increased HSAF production compared with that of the Δ*rpfG* mutant strain. Importantly, we found that the concentration of the Δ*rpfG* mutant did not significantly change c-di-GMP production compared to that of the wild-type strain in the HSAF-production medium (1/10 TSB). These data demonstrated that the regulatory activity of RpfG does not depend on its PDE enzymatic activity. This is the first report showing that a PDE does not depend on its c-di-GMP-degrading activity to regulate a downstream pathway. Thus, we wondered whether RpfG regulates HSAF synthesis through interactions with other proteins in *L. enzymogenes*.

Bioinformatics predictions found that RpfG may interact with three HyTCS proteins (HtsH1, HtsH2 and HtsH3). Then, we used pull-down and SPR to demonstrate the binding events between the RpfG and HtsH1, HtsH2 or HtsH3 proteins. Notably, RpfG and HtsH1, HtsH2 or HtsH3 interactions affect the PDE activity of RpfG. However, how RpfG affects HtsH1, HtsH2 or HtsH3 remains unknown. We speculate that RpfG may affect HtsH1, HtsH2 or HtsH3 autophosphorylation. However, we cannot obtain the full-length HtsH1, HtsH2 and HtsH3 proteins, so further clarification of these possible mechanisms will help elucidate the mechanism underlying the RpfG interaction with HtsH1, HtsH2 or HtsH3. On the other hand, we found that *htsH1*, *htsH2* and *htsH3* likely constitute a single transcription unit. HyTCS-based regulation may be crucial for responding to environmental changes and finely tuning gene expression (Borland *et al.*, 2016; Cui *et al.*, 2011; Parkinson *et al.*, 2015; Chambonnier *et al.*, 2016). However, the biological function of three consecutive HyTCS proteins has not been reported in bacteria.

In this study, we found that the in-frame deletion of the *htsH1*, *htsH2* and *htsH3* coding sequences significantly decreased HSAF production. Thus, we speculate that RpfG may interact with three HyTCS proteins to coordinate HSAF production in *L. enzymogenes*. To test this hypothesis, we performed an *E. coli*-based one-hybrid assay and EMSA. The results demonstrated that HtsH1, HtsH2 and HtsH3 could directly target the promoters of HSAF biosynthesis genes. We further analysed the transcription level of HSAF biosynthesis-related genes in *htsH1*, *htsH2* and *htsH3* mutants. Knockout of *htsH1*, *htsH2* and *htsH3* significantly reduced the transcription level of HSAF biosynthesis genes. These results suggest that HtsH1, HtsH2 and HtsH3 can directly regulate HSAF biosynthesis gene expression and increase HSAF production in *L. enzymogenes*.

Phosphorylation of TCS is critical for regulating the expression of downstream genes (Deng *et al.*, 2018; Cheng *et al.*, 2019; Wang *et al.*, 2017). However, whether HtsH1, HtsH2 and HtsH3, as a group of HyTCS proteins, regulate the expression of HSAF biosynthesis genes depends on their phosphorylation. We used Mn^2+^-Phos-tag SDS-PAGE and EMSA to show that HtsH1, HtsH2 and HtsH3 target p*lafB* depending on their phosphorylation activity. In this study, we report for the first time the biological functions of the three HyTCS proteins HtsH1, HtsH2 and HtsH3.

One of the notable results of this study is that RpfG, and HtsH1, HtsH2 and HtsH3 regulatory patterns are conserved mechanisms in *Lysobacter* and *Xanthomonas*. To our knowledge, RpfG represents a unique example of a c-di-GMP metabolic enzyme that directly interacts with three HyTCS proteins (HtsH1, HtsH2 and HtsH3) to regulate HSAF biosynthesis.

## Materials and methods

### Bacterial strains, plasmids, and growth conditions

The strains and plasmids used in this study are shown in Supplementary Table S1. *E. coli* strains were grown in Luria-Bertani medium (10 g/L tryptone, 5 g/L yeast extract, 10 g/L NaCl, pH 7.0) at 37°C. *L. enzymogenes* strains were grown at 28°C in Luria-Bertani medium and 10% TSB. For the preparation of culture media, tryptone, peptone, beef extract, and yeast extract were purchased from Sangon Biotech (Shanghai, China). When required, antibiotics were added (30 μg/mL kanamycin sulfate, 50 μg/mL gentamycin) to the *E. coli* or *L. enzymogenes* cultures. The bacterial growth in liquid medium was determined by measuring the optical density at 600 nm (OD600) using a Bioscreen-C Automated Growth Curves Analysis System (OY Growth Curves FP-1100-C, Helsinki, Finland).

### Site-directed mutagenesis

Site-directed mutagenesis and essentiality testing were performed as described previously (Li *et al.*, 2020a). To obtain the RpfG mutant proteins and *rpfG* site-directed mutant strains, plasmids harbouring mutations in *rpfG* were constructed. For example, to obtain the H190A mutation in RpfG, approximately 500-bp DNA fragments flanking the *rpfG* gene were amplified with *Pfu* DNA polymerase using *L. enzymogenes* genomic DNA as template and either MBP-*rpfG* P1 and *rpfG* H190A P1 (for the Up *rpfG* H190A mutant), or *rpfG* H190A P1 and MBP-*rpfG* P2 (for the Down *rpfG* H190A mutant) as the primers (Supplementary Table S2). The fragments were connected by overlap PCR using the primers MBP-*rpfG* P1 and MBP-*rpfG* P2. The fused fragment was digested with *Bam*H I and *Hin*dIII and inserted into pMAl-p2x to obtain the plasmid pMAl-*rpfG* H190A. The other four site-directed mutant plasmids (D191A, G253A, Y254A, and P255A) were constructed using a similar method.

### Protein expression and purification

Protein expression and purification were performed as described previously (Li *et al.*, 2017). To clone the *rpfG* gene, genomic DNA extracted from *L. enzymogenes* was used for PCR amplification using *Pfu* DNA polymerase, and the primers are listed in Supplementary Table S2. The PCR products were inserted into pMAl-p2x to produce the plasmids pMAl-*rpfG*. The *rpfG* gene was verified by nucleotide sequencing by Genscript (Nanjing, Jiangsu, China). Le *rpfG* and *rpfG* site-directed mutants with a vector-encoded maltose binding protein in the N-terminus were expressed in *E. coli* BL21 (DE3) and purified with dextrin sepharose high performance (Qiagen, Chatsworth, CA, USA) using an affinity column (Qiagen). The protein purity was monitored by SDS-PAGE. His_6_-tagged protein expression and purification were performed as described previously (Li *et al.*, 2020a; Li *et al.*, 2020b; Li *et al.*, 2017).

### PDE activity assays *in vitro*

The PDE activity assay was performed essentially as described earlier (Xu *et al.*, 2018). Briefly, 2 μM MBP-RpfG or its derivatives were tested in buffer containing 60 mM Tris-HCl (pH 7.6), 50 mM NaCl, 10 mM MnCl_2_ and 10 mM MgCl_2_. The reaction was started by the addition of 100 μM c-di-GMP. All reaction mixtures were incubated at 28°C for 5 to 60 min, followed by boiling for 10 min to stop the reaction. The samples were filtered through a 0.2 μM pore size cellulose-acetate filter, and 20 μl of each sample was loaded onto a reverse-phase C18 column and separated by HPLC. The separation protocol involved two mobile phases, 100 mM KH_2_PO_4_ plus 4 mM tetrabutylammonium sulfate (A) and 75% A + 25% methanol (B).

### C-di-GMP extraction and quantification

C-di-GMP extraction and quantification were performed as described previously (Xu *et al.*, 2018). Cultures were grown in 1/10 TSB at 28°C until the OD600 reached 1.5 based on the growth curve. Cells from 2 mL of the culture were harvested for protein quantification by BCA (TransGen, China). Cells from 8 mL of culture were used for c-di-GMP extraction using 0.6 M HClO_4_ and 2.5 M K_2_CO_3_. The samples were subjected to 0.22 µm Mini-Star filtration, and the filtrate was concentrated to 100 µL for liquid chromatography-tandem mass spectrometry (LC-MS/MS) analysis on an AB SCIEX QTRAP 6500 LC-MS/MS system (AB SCIEX, USA). The separation protocol involved two mobile phases, buffer A: 100 mM ammonium acetate plus 0.1% acetic acid and buffer B: 100% methanol. The gradient system was from 90% buffer A and 10% buffer B to 20% buffer A and 80% buffer B. The running time was 9 min, and the flow rate was 0.3 mL/min.

### Gene deletion and complementation

The in-frame deletions in *L. enzymogenes* OH11 were generated via double-crossover homologous recombination as described previously (Li *et al.*, 2020a; Xu *et al.*, 2018) using the primers listed in Supplementary Table S2. In brief, the flanking regions of each gene were PCR-amplified and cloned into the suicide vector pEX18Gm (Supplementary Table S1). The deletion constructs were transformed into the wild-type strain by electroporation, and gentamycin was used to select for integration of the nonreplicating plasmid into the recipient chromosome. A single-crossover integrant colony was spread on LB medium without gentamycin and incubated at 28°C for 3 days, and after appropriate dilution the culture was spread on LB plates containing 15% sucrose. Colonies sensitive to gentamycin were screened by PCR using the primers listed in Supplementary Table S2, and the gene deletion strains were obtained.

For gene complementation constructs, DNA fragments containing the full-length genes along with their promoters were PCR amplified and cloned into the versatile plasmid pBBR1MCS5 (Kovach *et al.*, 1995). The resulting plasmids were transferred into the *L. enzymogenes* strain by electroporation and the transformants were selected on LB plates containing Gm.

### RNA-Seq

The RNA-Seq assay was performed as described previously (Yang *et al.*, 2017; Zhou *et al.*, 2017). Briefly, the wild-type, Δ*htsH1*, Δ*htsH2*, Δ*htsH3*, Δ*htsH12*, Δ*htsH13*, Δ*htsH23* and Δ*htsH123* mutant strains were grown in 1/10 TSB medium at 28°C, and their cells were collected when the OD600 reached 1.0 based on the growth curve. The collected cells were used for RNA extraction by the TRIzol-based method (Life Technologies, CA, USA), and RNA degradation and contamination were monitored on 1% agarose gels. Then, clustering and sequencing were performed by Genedenovo Biotechnology Co., Ltd (Guangzhou, Guangdong, China). To analyse the DEGs between the wild-type, Δ*htsH1*, Δ*htsH2*, Δ*htsH3*, Δ*htsH12*, Δ*htsH13*, Δ*htsH23* and Δ*htsH123* mutant strains, the gene expression levels were further normalized using the fragments per kilobase of transcript per million (FPKM) mapped reads method to eliminate the influence of different gene lengths and amounts of sequencing data on the calculation of gene expression. The edgeR package (http://www.r-project.org/) was used to determine DEGs across samples with fold changes ≥ 2 and a false discovery rate-adjusted P (q value) < 0.05. DEGs were then subjected to enrichment analysis of GO functions and KEGG pathways, and q values were corrected using < 0.05 as the threshold.

### Quantitative real-time PCR

Quantitative real-time PCR was carried out according to previous studies (Cui *et al.*, 2018). The bacterial cells were collected when the cellular optical density (OD600) reached 1.0 in 10% TSB. Total RNA was extracted using a TRIzol-based method (Life Technologies, CA, USA). RNA quality control was performed via several steps: (1) the degree of RNA degradation and potential contamination were monitored on 1% agarose gels; (2) the RNA purity (OD260/OD280, OD260/OD230) was checked using a NanoPhotometer® spectrophotometer (IMPLEN, CA, USA); and (3) the RNA integrity was measured using a Bioanalyser 2100 (Agilent, Santa Clara, CA, USA). The primers used in this assay are listed in Supplementary Table S2. cDNA was then synthesized from each RNA sample (400 ng) using the TransScript^®^ All-in-One First-Strand cDNA Synthesis SuperMix for qPCR (One-Step gDNA Removal) Kit (TransGen Biotech, Beijing, China) according to the manufacturer’s instructions. qRT-PCR was performed using TransStart Top Green qPCR SuperMix (TransGen Biotech) on a QuantStudio TM 6 Flex Real-Time PCR System (Applied Biosystems, Foster City, CA, USA) with the following thermal cycling parameters: denaturation at 94°C for 30 s, followed by 40 cycles of 94°C for 5 s and 60°C for 34 s. Gene expression analyses were performed using the 2^−ΔΔCT^ method with 16S rDNA as the endogenous control, and the expression level in the wild type was set to a value of 1. The experiments were performed three times, and three replicates were examined in each run.

### Pull-down assay

The assay was performed as described previously (Xu *et al.*, 2018). Briefly, the purified proteins were used to perform the pull-down assay in a reaction system comprising 800 µL PBS buffer, 5 µM (final concentration) MBP-RpfG and HtsH1C-Flag-His, HtsH2C-HA-His or HtsH3C-Myc-His proteins, and 50 µL Dextrin Sepharose High Performance agarose (Sigma-Aldrich, St. Louis, MO, USA). All samples were incubated at 4°C overnight. The agarose was collected by centrifugation and washed 10 times with PBS containing 1% Triton X-100 to remove non-specifically bound proteins. The MBP-bead-captured proteins were eluted by boiling in 6× SDS loading dye for 10 min. These samples were subjected to SDS-PAGE and Western blotting. Protein detection involved the use of MBP- (ab49923), Flag- (ab1162), HA- (ab187915), Myc- (ab32072) and His- (ab18184) specific antibodies obtained from Abcam, UK.

### Phosphorylation analysis through Phos-tag gel

The purified HtsH1C-Flag-His, HtsH2C-HA-His or HtsH3C-Myc-His proteins (100 ng) were incubated with CIAP (Solarbio, Beijing, China) at 28°C for 60 min, and resolved on 8% SDS-PAGE prepared with 50 µM acrylamide-dependent Phos-tag ligand and 100 µM MnCl_2_ as previously described (Yin *et al.*, 2020). Gel electrophoresis was performed with a constant voltage of 80 V for 3-6 h. Before transferring, gels were equilibrated in transfer buffer with 5 mM EDTA for 20 min two times, followed by transfer buffer without EDTA for another 20 min. Protein transfer from the Mn^2+^ phos-tag acrylamide (APExBIO, Houston, USA) gel to the PVDF membrane (Millipore, Massachusetts, USA) was performed for ∼24 h at 80 V at 4°C, and then the membrane was analysed by Western blotting using the anti-His antibody.

### Bacterial one-hybrid assays

Bacterial one-hybrid assays were performed as previously described (Xu *et al.*, 2016; Wang *et al.*, 2018). In brief, the bacterial one-hybrid reporter system contains three components: the plasmids pBXcmT and pTRG, which are used to clone the target DNA and to express the target protein, respectively, and the *E. coli* XL1-Blue MRF′ kan strain, which is the host strain for the propagation of the pBXcmT and pTRG recombinants (Guo *et al.*, 2009). In this study, the promoter of the HSAF biosynthesis gene (p*lafB*) was cloned into pBXcmT to generate the recombinant vector pBXcmT-p*lafB*. Similarly, the coding regions of Le *htsH1*, Le *htsH2*, and Le *htsH3* were cloned into pTRG to create the final constructs pTRG-*htsH1*, pTRG-*htsH2*, and pTRG-*htsH3*, respectively. The two recombinant vectors were transformed into the XL1-Blue MRF′ kan strain. If direct physical binding occurred between HtsH1, HtsH2 or HtsH3 and p*lafB*, the positive-transformant *E. coli* strain containing both pBXcmT-p*lafB* and pTRG-HtsHs would grow well on selective medium, that is, minimal medium containing 5 mM 3-amino-1,2,4-triazole, 8 μg/mL streptomycin, 12.5 μg/mL tetracycline, 34 μg/mL chloramphenicol and 30 μg/mL kanamycin. Furthermore, cotransformants containing pBX-R2031/pTRG-R3133 served as a positive control (Guo *et al.*, 2009), and cotransformants containing either empty pTRG or pBXcmT-p*lafB* were used as negative controls. All cotransformants were spotted onto selective medium, grown at 28°C for 3-4 days, and then photographed.

### Electrophoretic mobility gel shift assays (EMSAs)

EMSA was performed as previously described (Hirakawa *et al.*, 2015; Shao *et al.*, 2018). For HtsH1, HtsH2 or HtsH3 gel shift assays, we used DNA fragments that included p*lafB* as a probe. The probe DNA (50 ng) was mixed with protein in a 20 μL reaction mixture containing 10 mM Tris-HCl (pH 7.5), 50 mM KCl, 1 mM dithiothreitol, and 0.4% glycerol. After incubation for 30 min at 28°C, samples were electrophoresed on a 5% nondenaturing acrylamide gel in 0.5× TBE buffer at 4°C. The gel was soaked in 10,000- fold-diluted SYBR Green I nucleic acid dye (Sangon Biotech, Shanghai, China), and the DNA was visualized at 300 nm.

### HSAF extraction and quantification

HSAF was extracted from 50 mL *L. enzymogenes* cultures grown in 10% TSB for 48 h at 28°C with shaking (at 180 rpm). HSAF was detected via HPLC and quantified per unit of OD600 as described previously (Qian *et al.*, 2013; Wang *et al.*, 2014; Xu *et al.*, 2018). Three biological replicates were used, and each was examined with three technical replicates.

### Statistical analyses

The experimental datasets were subjected to analyses of variance using GraphPad Prism 7.0. The significance of the treatment effects was determined by the F value (*P* = 0.05). If a significant F value was obtained, separation of means was accomplished by Fisher’s protected least significant difference at *P* ≤ 0.05.

## Acknowledgments

The authors thank Haihong Wang for helpful suggestions on the manuscript. It is also a great pleasure to thank Cunfa Xu at the Central Laboratory of Jiangsu Academy of Agricultural Sciences for SPR technical support.

## Additional information

### Funding

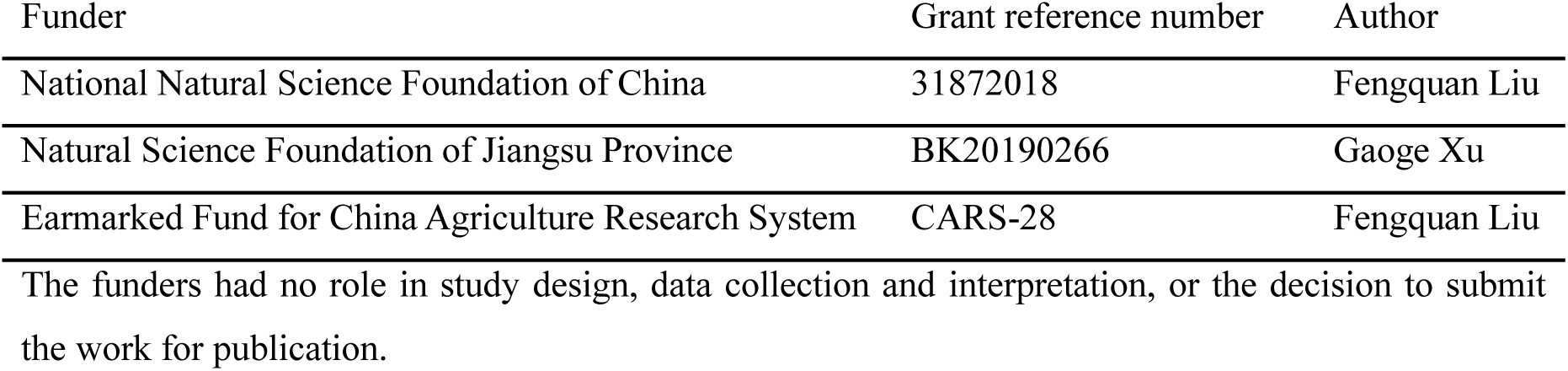

### Author contributions

K.L. and F.L. conceived and designed the experiments. K.L., G.X., B.W., and G.W. performed the experiments. K.L. analysed the data and prepared the figures. K.L. and F.L. wrote the manuscript draft. F.L. revised the manuscript. All authors read and approved the final manuscript.

## Additional files

### Supplementary files

Supplementary file 1. Key supplementary table.

### Data availability

The data that support the findings of this study are openly available in GenBank at https://www.ncbi.nlm.nih.gov/nuccore/, accession numbers RCTY01000033 (*Lysobacter enzymogenes* strain OH11 scffold34, whole genome shotgun sequence; Le4727/Le RpfG, locus tag = D9T17_13845), RCTY01000055 (*Lysobacter enzymogenes* strain OH11 scffold56, whole genome shotgun sequence, Le3071/Le HtsH1, locus tag = D9T17_21400)), RCTY01000054 (*Lysobacter enzymogenes* strain OH11 scffold55, whole genome shotgun sequence, Le3072/Le HtsH2, locus tag = D9T17_21390), RCTY01000054 (*Lysobacter enzymogenes* strain OH11 scffold55, whole genome shotgun sequence, Le3073/Le HtsH3, locus tag = D9T17_21385).

## FIGURE LEGENDS

**Figure 3—figure supplement 1.**
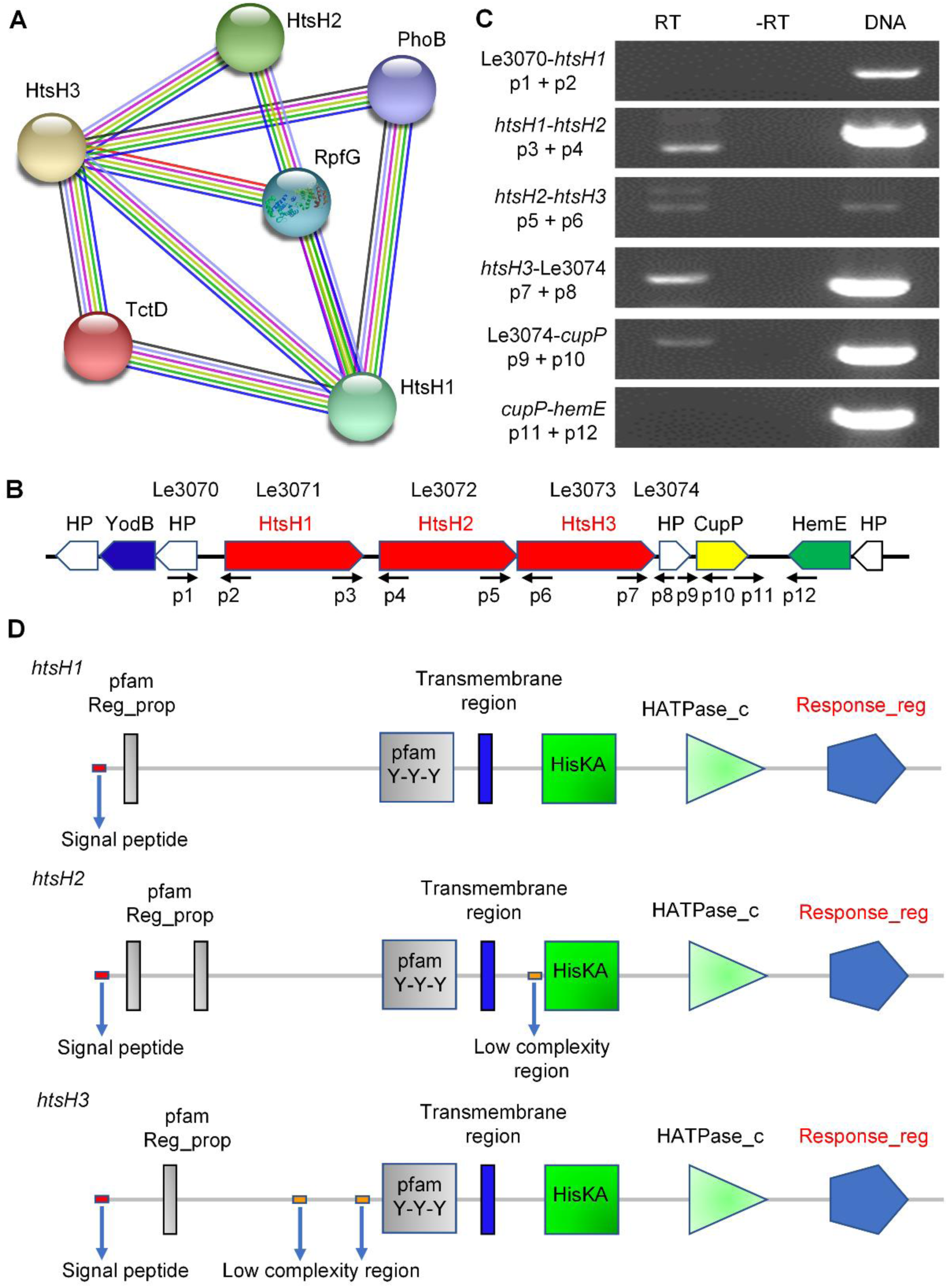
*htsH1*, *htsH2* and *htsH3* are located in an operon. **(A)** Bioinformatics predictions that RpfG may interact with HtsH1, HtsH2 and HtsH3. **(B)** Genomic localization of *htsH1*, *htsH2* and *htsH3*. Arrows indicate open reading frame regions of the genes and their transcriptional directions. Gene names are listed above, and primers used to verify operon structures by RT-PCR are indicated below. The primers used are listed in Supplementary Table S2. HP: hypothetical protein; YodB: cytochrome B561 (yodB); HP: hypothetical protein; HtsH1: Hybrid two-component systems protein; HtsH2: Hybrid two-component systems protein; HtsH3: Hybrid two-component systems protein; HP: hypothetical protein; CupP: Cupin 2 conserved barrel domain protein; HemE: uroporphyrinogen decarboxylase (hemE); HP: hypothetical protein. **(C)** Verification of operon organization by RT-PCR. The cDNA was reverse-transcribed with random primers using total RNA from *L. enzymogenes* grown in 10% TSB medium at 28°C until the OD600 reached 1.0. PCR fragment lengths are listed on the right. RT represents amplification using cDNA transcribed from RNA as template; -RT represents the negative control, in which reverse transcriptase was absent during cDNA synthesis; DNA represents the positive control using DNA as the PCR template. **(D)** Bioinformatics analyses of the domain organization of HtsH1, HtsH2 and HtsH3 that belong to HyTCS.

**Figure 3—figure supplement 2.**
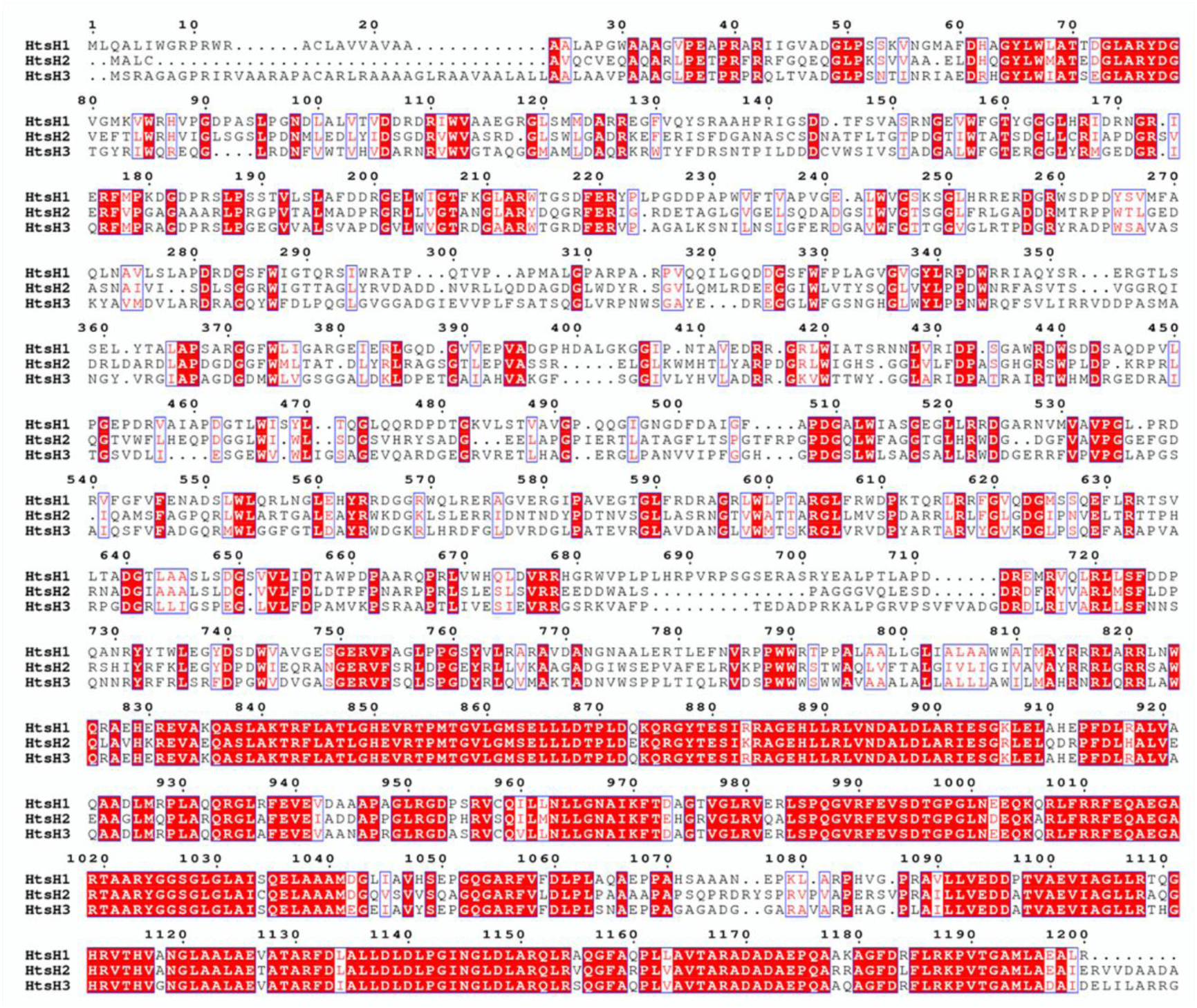
Identification and sequence characterization of HtsH1, HtsH2 and HtsH3 in *L. enzymogenes*. Alignment of *L. enzymogenes* HtsH1, HtsH2 and HtsH3. The alignment was performed with Clustal W based on identical residues.

**Figure 3—figure supplement 3.**
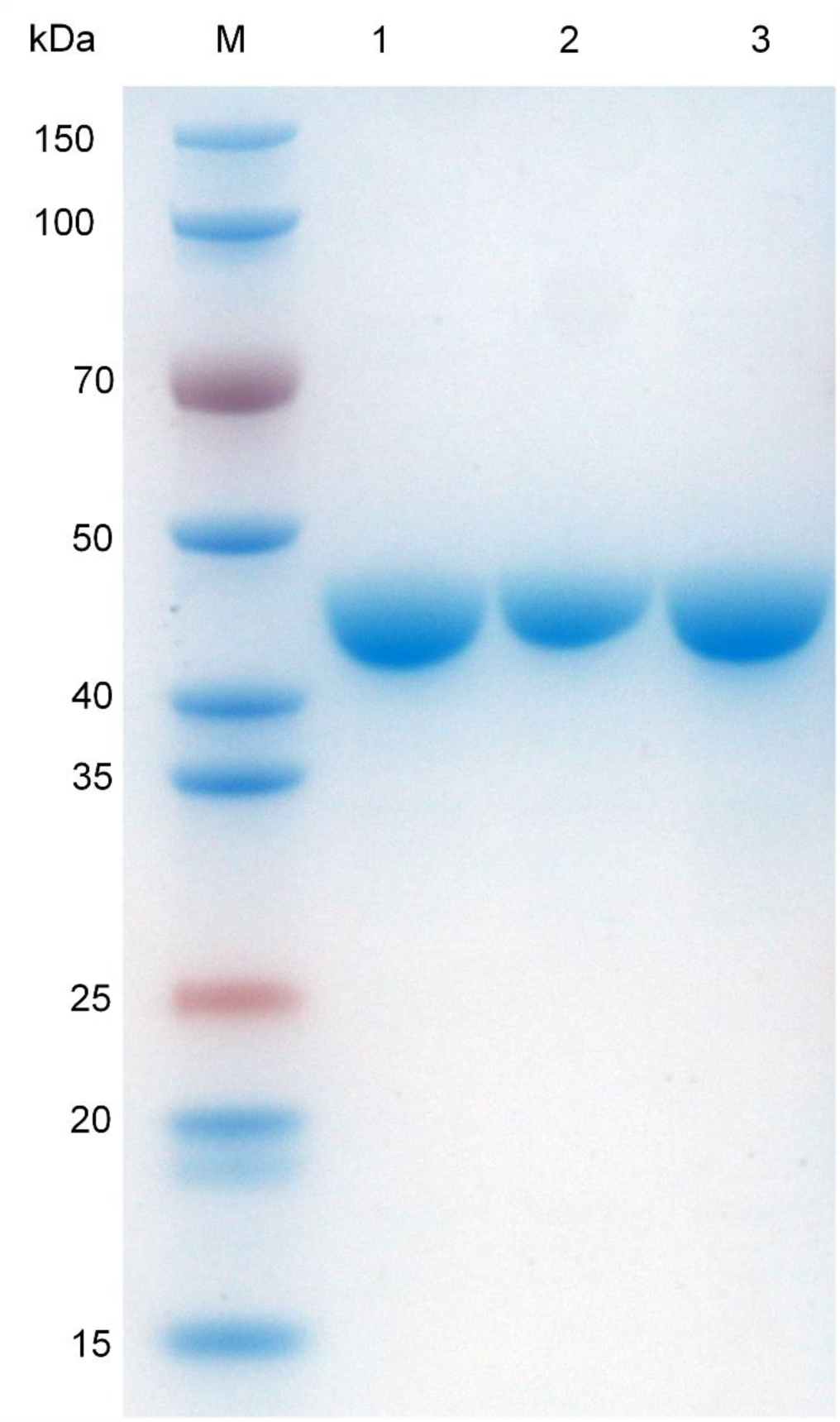
The purified cytoplasmic fragments of HtsH1, HtsH2 and HtsH3 were analysed by 12% SDS-PAGE. Lane M, molecular mass markers; lane 1, HtsH1C-Flag-His protein; lane 2, HtsH2C-HA-His protein; lane 3, HtsH3C-Myc-His protein.

**Figure 3—figure supplement 4.**
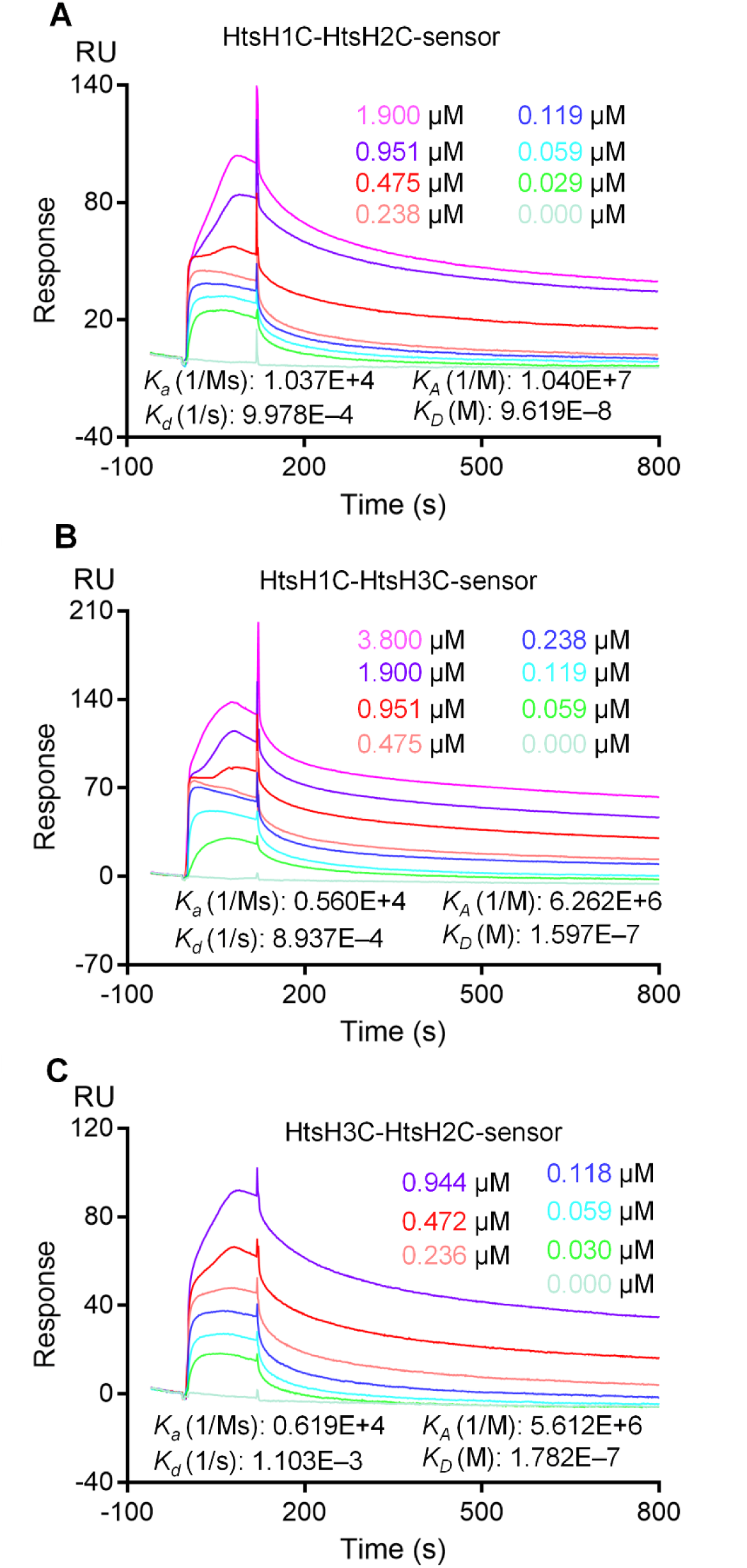
HtsH1, HtsH2 and HtsH3 exist as a complex in *L. enzymogenes*. **(A)** SPR showing that HtsH1C-Flag-His forms a complex with HtsH2C- HA-His with K_D_ = 0.09619 nM. **(B)** SPR showing that HtsH1C-Flag-His forms a complex with HtsH3C-Myc-His with K_D_ = 0.1597 nM. **(C)** SPR showing that HtsH2C-HA-His forms a complex with HtsH3C-Myc-His with K_D_ = 0.1782 nM.

**Figure 6—figure supplement 1.**
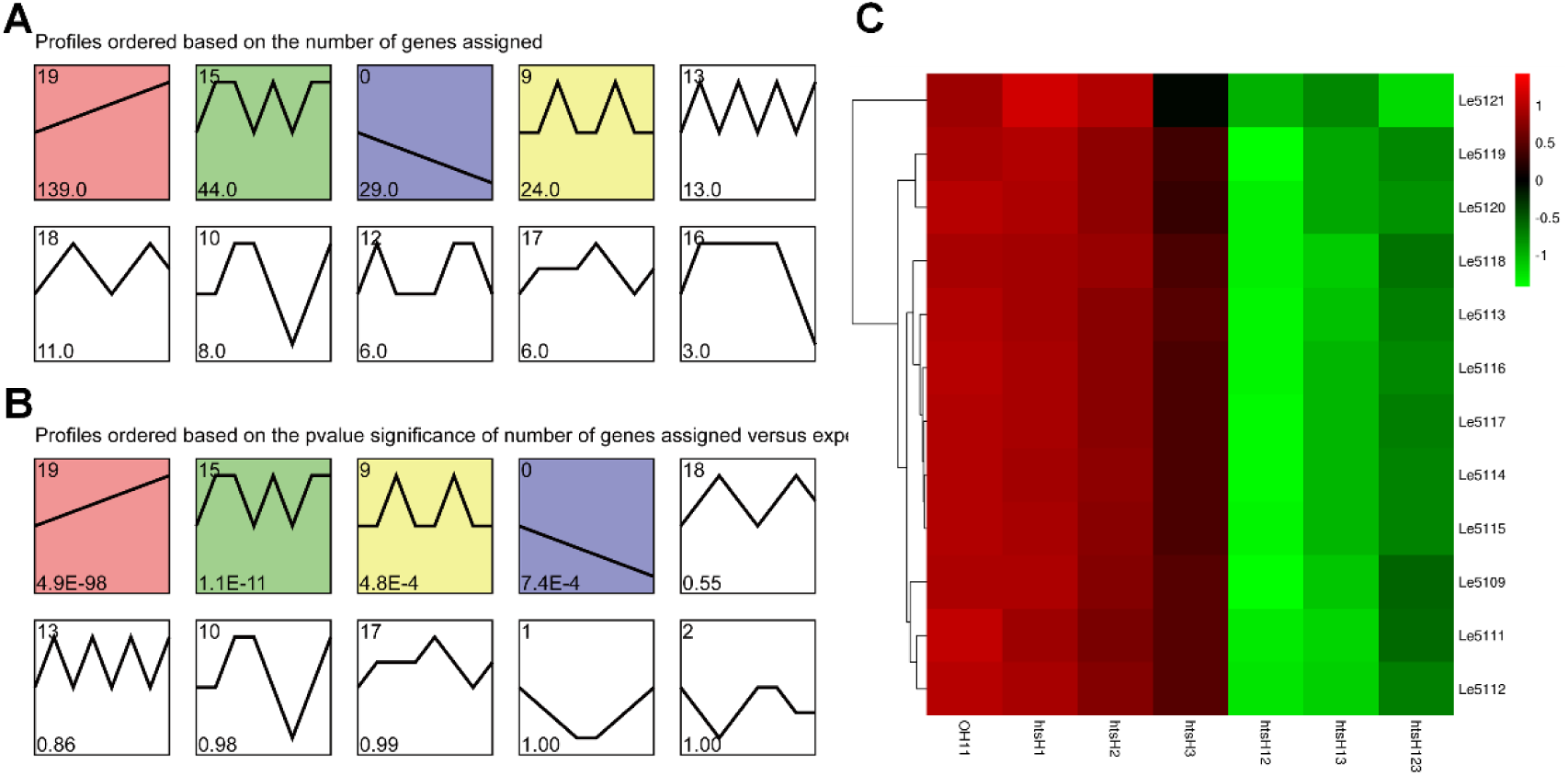
Transcriptional analysis of HSAF biosynthesis genes in *L. enzymogenes*. **(A-B)** Trend analysis of differential gene expression in the *htsHs* mutants (Δ*htsH1*, Δ*htsH2*, Δ*htsH3*, Δ*htsH12*, Δ*htsH13*, Δ*htsH23* and Δ*htsH123*). **(C)** Hierarchical cluster analysis applied to the 12 DEGs in the HSAF biosynthesis gene cluster in different mutant backgrounds.

**Figure 8—figure supplement 1.**
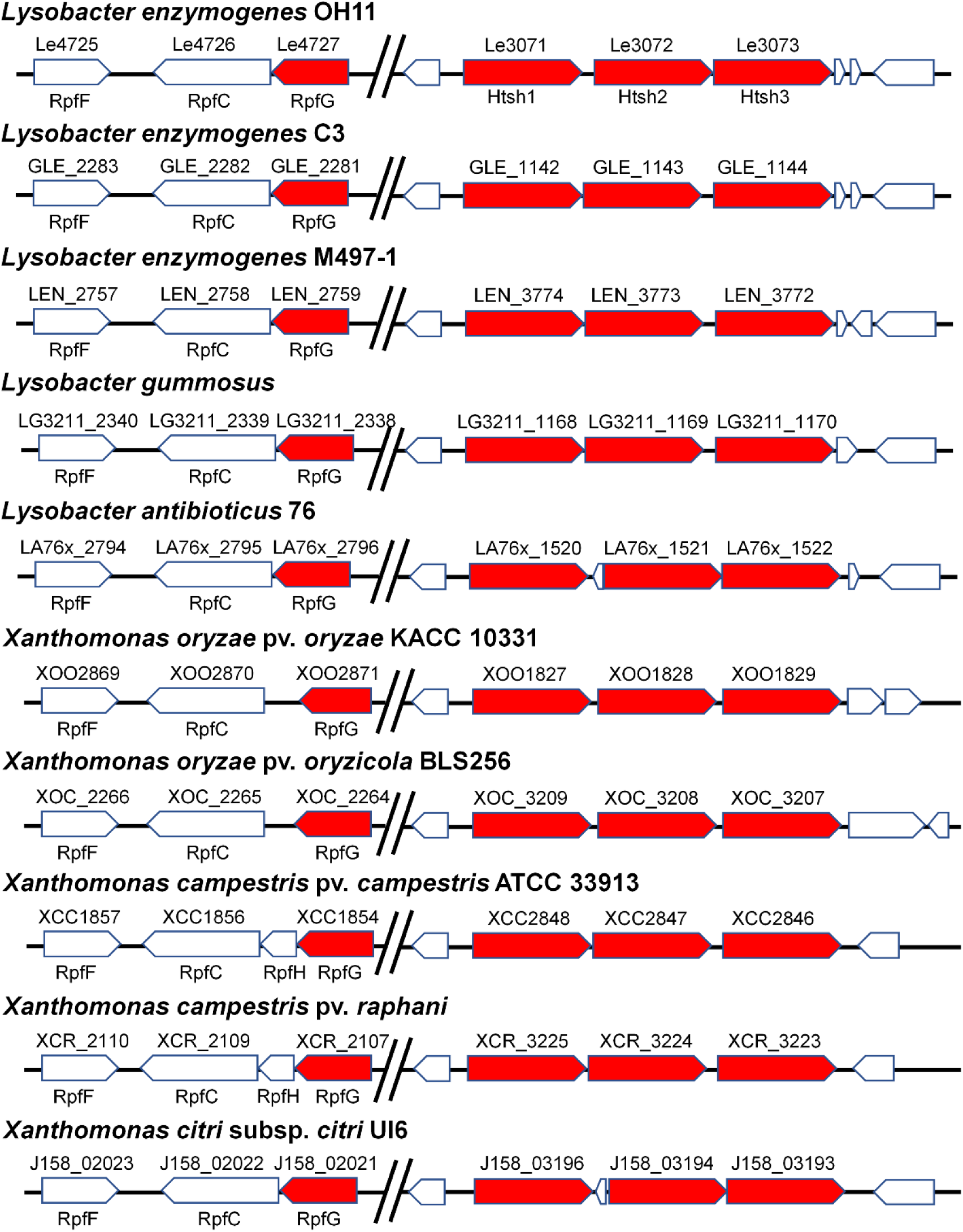
Conservation of the key genes for RpfG, and HtsH1, HtsH2 and HtsH3-dependent regulatory patterns in the genomes of different bacteria. Genomic organization of the genes and homology analysis of the products. All sequences were retrieved from NCBI Microbial Genome Resources. All amino acid sequences were downloaded from the microbial genome sequence database of NCBI. Position-specific Iterated BLAST (PSI-BLAST) was used for homology analysis.

**Supplementary Table S1.** Bacterial strains and plasmids used in this study.

**Supplementary Table S2.** Sequences of the PCR primers used in this work.

**Supplementary Table S3.** List of genes differentially expressed in the *htsH1*, *htsH2*, and *htsH3* mutants compared to the wild-type strain (log2 fold change ≥ 1).

## References

Andrade MO, Alegria MC, Guzzo CR, Docena C, Rosa MC, Ramos CH, Farah CS. 2006. The HD-GYP domain of RpfG mediates a direct linkage between the Rpf quorum-sensing pathway and a subset of diguanylate cyclase proteins in the phytopathogen Xanthomonas axonopodis pv citri. Molecular microbiology 62:537–551. DOI: https://doi.org/10.1111/j.1365-2958.2006.05386.x, PMID: 17020586

Azcarate-Peril MA, McAuliffe O, Altermann E, Lick S, Russell WM, Klaenhammer TR. 2005. Microarray analysis of a two-component regulatory system involved in acid resistance and proteolytic activity in Lactobacillus acidophilus. Applied and environmental microbiology 71:5794–5804. DOI: https://doi.org/10.1128/AEM.71.10.5794-5804.2005, PMID: 16204490

Barber CE, Tang JL, Feng JX, Pan MQ, Wilson TJ, Slater H, Dow JM, Williams P, Daniels MJ. 1997. A novel regulatory system required for pathogenicity of Xanthomonas campestris is mediated by a small diffusible signal molecule. Molecular microbiology 24:555–566. DOI: https://doi.org/10.1046/j.1365-2958.1997.3721736.x, PMID: 9179849

Bellini D, Caly DL, McCarthy Y, Bumann M, An SQ, Dow JM, Ryan RP, Walsh MA. 2014. Crystal structure of an HD-GYP domain cyclic-di-GMP phosphodiesterase reveals an enzyme with a novel trinuclear catalytic iron centre. Molecular microbiology 91:26–38. DOI: https://doi.org/10.1111/mmi.12447, PMID: 24176013

Borland S, Prigent-Combaret C, Wisniewski-Dye F. 2016. Bacterial hybrid histidine kinases in plant-bacteria interactions. Microbiology 162:1715–1734. DOI: https://doi.org/10.1099/mic.0.000370, PMID: 27609064

Chambonnier G, Roux L, Redelberger D, Fadel F, Filloux A, Sivaneson M, de Bentzmann S, Bordi C. 2016. The Hybrid Histidine Kinase LadS Forms a Multicomponent Signal Transduction System with the GacS/GacA Two-Component System in Pseudomonas aeruginosa. PLoS genetics 12:e1006032. DOI: https://doi.org/10.1371/journal.pgen.1006032, PMID: 27176226

Chatterjee S, Sonti RV. 2002. rpfF mutants of Xanthomonas oryzae pv. oryzae are deficient for virulence and growth under low iron conditions. Molecular plant-microbe interactions : MPMI 15:463–471. DOI: https://doi.org/10.1094/MPMI.2002.15.5.463, PMID: 12036277

Chen Y, Xia J, Su Z, Xu G, Gomelsky M, Qian G, Liu F. 2017. Lysobacter PilR, the Regulator of Type IV Pilus Synthesis, Controls Antifungal Antibiotic Production via a Cyclic di-GMP Pathway. Applied and environmental microbiology 83:e03397–03316. DOI: https://doi.org/10.1128/aem.03397-16, PMID: 28087536

Cheng ST, Wang FF, Qian W. 2019. Cyclic-di-GMP binds to histidine kinase RavS to control RavS-RavR phosphotransfer and regulates the bacterial lifestyle transition between virulence and swimming. PLoS pathogens 15:e1007952. DOI: https://doi.org/10.1371/journal.ppat.1007952, PMID: 31408509

Cui C, Yang C, Song S, Fu S, Sun X, Yang L, He F, Zhang LH, Zhang Y, Deng Y. 2018. A novel two-component system modulates quorum sensing and pathogenicity in Burkholderia cenocepacia. Molecular microbiology 108:32–44. DOI: https://doi.org/10.1111/mmi.13915, PMID: 29363827

Cui Y, Tu R, Wu L, Hong Y, Chen S. 2011. A hybrid two-component system protein from Azospirillum brasilense Sp7 was involved in chemotaxis. Microbiol Res 166:458–467. DOI: https://doi.org/10.1016/j.micres.2010.08.006, PMID: 20869221

Deng CY, Zhang H, Wu Y, Ding LL, Pan Y, Sun ST, Li YJ, Wang L, Qian W. 2018. Proteolysis of histidine kinase VgrS inhibits its autophosphorylation and promotes osmostress resistance in Xanthomonas campestris. Nature communications 9:4791. DOI: https://doi.org/10.1038/s41467-018-07228-4, PMID: 30442885

Deng Y, Wu J, Tao F, Zhang LH. 2011. Listening to a new language: DSF-based quorum sensing in Gram-negative bacteria. Chemical reviews 111:160–173. DOI: https://doi.org/10.1021/cr100354f, PMID: 21166386

Galperin MY, Natale DA, Aravind L, Koonin EV. 1999. A specialized version of the HD hydrolase domain implicated in signal transduction. Journal of molecular microbiology and biotechnology 1:303–305. PMID: 10943560

Galperin MY, Nikolskaya AN, Koonin EV. 2001. Novel domains of the prokaryotic two-component signal transduction systems. FEMS Microbiol Lett 203:11–21. DOI: https://doi.org/10.1111/j.1574-6968.2001.tb10814.x, PMID: 11557134

Goodman AL, Merighi M, Hyodo M, Ventre I, Filloux A, Lory S. 2009. Direct interaction between sensor kinase proteins mediates acute and chronic disease phenotypes in a bacterial pathogen. Genes & development 23:249–259. DOI: https://doi.org/10.1101/gad.1739009, PMID: 19171785

Guo M, Feng H, Zhang J, Wang W, Wang Y, Li Y, Gao C, Chen H, Feng Y, He ZG. 2009. Dissecting transcription regulatory pathways through a new bacterial one-hybrid reporter system. Genome research 19:1301–1308. DOI: https://doi.org/10.1101/gr.086595.108, PMID: 19228590

Han Y, Wang Y, Tombosa S, Wright S, Huffman J, Yuen G, Qian G, Liu F, Shen Y, Du L. 2015. Identification of a small molecule signaling factor that regulates the biosynthesis of the antifungal polycyclic tetramate macrolactam HSAF in Lysobacter enzymogenes. Applied microbiology and biotechnology 99:801–811. DOI: https://doi.org/10.1007/s00253-014-6120-x, PMID: 25301587

He YW, Wang C, Zhou L, Song H, Dow JM, Zhang LH. 2006. Dual signaling functions of the hybrid sensor kinase RpfC of Xanthomonas campestris involve either phosphorelay or receiver domain-protein interaction. The Journal of biological chemistry 281:33414–33421. DOI: https://doi.org/10.1074/jbc.M606571200, PMID: 16940295

Hirakawa H, Hirakawa Y, Greenberg EP, Harwood CS. 2015. BadR and BadM Proteins Transcriptionally Regulate Two Operons Needed for Anaerobic Benzoate Degradation by Rhodopseudomonas palustris. Applied and environmental microbiology 81:4253–4262. DOI: https://doi.org/10.1128/AEM.00377-15, PMID: 25888170

Kovach ME, Elzer PH, Hill DS, Robertson GT, Farris MA, Roop RM, 2nd, Peterson KM. 1995. Four new derivatives of the broad-host-range cloning vector pBBR1MCS, carrying different antibiotic-resistance cassettes. Gene 166:175–176. DOI: https://doi.org/10.1016/0378-1119(95)00584-1, PMID: 8529885

Li K-H, Yu Y-H, Dong H-J, Zhang W-B, Ma J-C, Wang H-H. 2017. Biological functions of ilvC in branched-chain fatty acid synthesis and diffusible signal factor family production in Xanthomonas campestris. Frontiers in microbiology 8:2486. DOI: https://doi.org/10.3389/fmicb.2017.02486, PMID: 29312195

Li K, Hou R, Xu H, Wu G, Qian G, Wang H, Liu F. 2020a. Two functional fatty acyl coenzyme A ligases affect free fatty acid metabolism to block biosynthesis of an antifungal antibiotic in Lysobacter enzymogenes. Applied and environmental microbiology 86:e00309–00320. DOI: https://doi.org/10.1128/aem.00309-20, PMID: 32144106

Li K, Wu G, Liao Y, Zeng Q, Wang H, Liu F. 2020b. RpoN1 and RpoN2 play different regulatory roles in virulence traits, flagellar biosynthesis, and basal metabolism in Xanthomonas campestris. Molecular plant pathology 21:907–922. DOI: https://doi.org/10.1111/mpp.12938, PMID: 32281725

Li S, Du L, Yuen G, Harris SD. 2006. Distinct ceramide synthases regulate polarized growth in the filamentous fungus Aspergillus nidulans. Molecular biology of the cell 17:1218–1227. DOI: https://doi.org/10.1091/mbc.e05-06-0533, PMID: 16394102

Li Y, Wang H, Liu Y, Jiao Y, Li S, Shen Y, Du L. 2018. Biosynthesis of the Polycyclic System in the Antifungal HSAF and Analogues from Lysobacter enzymogenes. Angew Chem Int Ed Engl 57:6221–6225. DOI: https://doi.org/10.1002/anie.201802488, PMID: 29573092

Lou L, Qian G, Xie Y, Hang J, Chen H, Zaleta-Rivera K, Li Y, Shen Y, Dussault PH, Liu F, Du L. 2011. Biosynthesis of HSAF, a tetramic acid-containing macrolactam from Lysobacter enzymogenes. Journal of the American Chemical Society 133:643–645. DOI: https://doi.org/10.1021/ja105732c, PMID: 21171605

Odhiambo BO, Xu G, Qian G, Liu F. 2017. Evidence of an Unidentified Extracellular Heat-Stable Factor Produced by Lysobacter enzymogenes (OH11) that Degrade Fusarium graminearum PH1 Hyphae. Current microbiology 74:437–448. DOI: https://doi.org/10.1007/s00284-017-1206-1, PMID: 28213660

Parkinson JS, Hazelbauer GL, Falke JJ. 2015. Signaling and sensory adaptation in Escherichia coli chemoreceptors: 2015 update. Trends in microbiology 23:257–266. DOI: https://doi.org/10.1016/j.tim.2015.03.003, PMID: 25834953

Qian G, Wang Y, Liu Y, Xu F, He YW, Du L, Venturi V, Fan J, Hu B, Liu F. 2013. Lysobacter enzymogenes uses two distinct cell-cell signaling systems for differential regulation of secondary-metabolite biosynthesis and colony morphology. Applied and environmental microbiology 79:6604–6616. DOI: https://doi.org/10.1128/AEM.01841-13, PMID: 23974132

Qian GL, Hu BS, Jiang YH, Liu FQ. 2009. Identification and Characterization of Lysobacter enzymogenes as a Biological Control Agent Against Some Fungal Pathogens. Agricultural Sciences in China 8:68–75. DOI: https://doi.org/Doi 10.1016/S1671-2927(09)60010-9

Ryan RP, McCarthy Y, Andrade M, Farah CS, Armitage JP, Dow JM. 2010. Cell-cell signal-dependent dynamic interactions between HD-GYP and GGDEF domain proteins mediate virulence in Xanthomonas campestris. Proceedings of the National Academy of Sciences of the United States of America 107:5989–5994. DOI: https://doi.org/10.1073/pnas.0912839107, PMID: 20231439

Shao X, Zhang X, Zhang Y, Zhu M, Yang P, Yuan J, Xie Y, Zhou T, Wang W, Chen S, Liang H, Deng X. 2018. RpoN-Dependent Direct Regulation of Quorum Sensing and the Type VI Secretion System in Pseudomonas aeruginosa PAO1. Journal of bacteriology 200:e00205–00218. DOI: https://doi.org/10.1128/JB.00205-18, PMID: 29760208

Su Z, Han S, Fu ZQ, Qian G, Liu F. 2018. Heat-Stable Antifungal Factor (HSAF) Biosynthesis in Lysobacter enzymogenes Is Controlled by the Interplay of Two Transcription Factors and a Diffusible Molecule. Applied and environmental microbiology 84: e01754–01717. DOI: https://doi.org/10.1128/AEM.01754-17, PMID: 29101199

Wang B, Wu G, Zhang Y, Qian G, Liu F. 2018. Dissecting the virulence-related functionality and cellular transcription mechanism of a conserved hypothetical protein in Xanthomonas oryzae pv. oryzae. Molecular plant pathology 19:1859–1872. DOI: https://doi.org/10.1111/mpp.12664, PMID: 29392817

Wang FF, Cheng ST, Wu Y, Ren BZ, Qian W. 2017. A Bacterial Receptor PcrK Senses the Plant Hormone Cytokinin to Promote Adaptation to Oxidative Stress. Cell reports 21:2940–2951. DOI: https://doi.org/10.1016/j.celrep.2017.11.017, PMID: 29212037

Wang Y, Zhao Y, Zhang J, Zhao Y, Shen Y, Su Z, Xu G, Du L, Huffman JM, Venturi V, Qian G, Liu F. 2014. Transcriptomic analysis reveals new regulatory roles of Clp signaling in secondary metabolite biosynthesis and surface motility in Lysobacter enzymogenes OH11. Applied microbiology and biotechnology 98:9009–9020. DOI: https://doi.org/10.1007/s00253-014-6072-1, PMID: 25236801

Xu G, Han S, Huo C, Chin KH, Chou SH, Gomelsky M, Qian G, Liu F. 2018. Signaling specificity in the c-di-GMP-dependent network regulating antibiotic synthesis in Lysobacter. Nucleic acids research 46:9276–9288. DOI: https://doi.org/10.1093/nar/gky803, PMID: 30202891

Xu H, Chen H, Shen Y, Du L, Chou SH, Liu H, Qian G, Liu F. 2016. Direct Regulation of Extracellular Chitinase Production by the Transcription Factor LeClp in Lysobacter enzymogenes OH11. Phytopathology 106:971–977. DOI: https://doi.org/10.1094/PHYTO-01-16-0001-R, PMID: 27385597

Yang C, Cui C, Ye Q, Kan J, Fu S, Song S, Huang Y, He F, Zhang LH, Jia Y, Gao YG, Harwood CS, Deng Y. 2017. Burkholderia cenocepacia integrates cis-2-dodecenoic acid and cyclic dimeric guanosine monophosphate signals to control virulence. Proceedings of the National Academy of Sciences of the United States of America 114:13006–13011. DOI: https://doi.org/10.1073/pnas.1709048114, PMID: 29158389

Yin Z, Feng W, Chen C, Xu J, Li Y, Yang L, Wang J, Liu X, Wang W, Gao C, Zhang H, Zheng X, Wang P, Zhang Z. 2020. Shedding light on autophagy coordinating with cell wall integrity signaling to govern pathogenicity of Magnaporthe oryzae. Autophagy 16:900–916. DOI: https://doi.org/10.1080/15548627.2019.1644075, PMID: 31313634

Yu F, Zaleta-Rivera K, Zhu X, Huffman J, Millet JC, Harris SD, Yuen G, Li XC, Du L. 2007. Structure and biosynthesis of heat-stable antifungal factor (HSAF), a broad-spectrum antimycotic with a novel mode of action. Antimicrobial agents and chemotherapy 51:64–72. DOI: https://doi.org/10.1128/aac.00931-06, PMID: 17074795

Zhao Y, Cheng C, Jiang T, Xu H, Chen Y, Ma Z, Qian G, Liu F. 2019. Control of Wheat Fusarium Head Blight by Heat-Stable Antifungal Factor (HSAF) from Lysobacter enzymogenes. Plant disease 103:1286–1292. DOI: https://doi.org/10.1094/PDIS-09-18-1517-RE, PMID: 30995421

Zhao Y, Qian G, Chen Y, Du L, Liu F. 2017. Transcriptional and Antagonistic Responses of Biocontrol Strain Lysobacter enzymogenes OH11 to the Plant Pathogenic Oomycete Pythium aphanidermatum. Frontiers in microbiology 8:1025. DOI: https://doi.org/10.3389/fmicb.2017.01025, PMID: 28634478

Zhou M, Shen D, Xu G, Liu F, Qian G. 2017. ChpA Controls Twitching Motility and Broadly Affects Gene Expression in the Biological Control Agent Lysobacter enzymogenes. Current microbiology 74:566–574. DOI: https://doi.org/10.1007/s00284-017-1202-5, PMID: 28258296

Zhou X, Qian G, Chen Y, Du L, Liu F, Yuen GY. 2015. PilG is Involved in the Regulation of Twitching Motility and Antifungal Antibiotic Biosynthesis in the Biological Control Agent Lysobacter enzymogenes. Phytopathology 105:1318–1324. DOI: https://doi.org/10.1094/PHYTO-12-14-0361-R, PMID: 26360465

